# Transitory Schwann Cell Precursor and hybrid states underpin melanoma therapy resistance and metastasis

**DOI:** 10.1101/2022.10.14.512297

**Authors:** Vishaka Gopalan, Chun Wai Wong, Rotem Leshem, Luke Owen, Tuulia Vallius, Yingxiao Shi, Yuhong Jiang, Eva Pérez-Guijarro, Emily Wu, Sung Chin, Jessica Ebersole, Cari Smith, Antonella Sassano, Maira Alves Constantino, Michael J. Haley, Ferenc Livak, R. Mark Simpson, Chi-Ping Day, Adam Hurlstone, Sridhar Hannenhalli, Glenn Merlino, Kerrie L. Marie

## Abstract

Melanoma plasticity, driven by phenotype state switching, underlies clinically relevant traits such as metastasis and therapy resistance. As melanoma progression is thought to recapitulate aspects of neural crest cell (NCC) development, understanding embryonic melanocyte specification and lineage fate decisions of closely related NCCs may illuminate the pathways co-opted during disease evolution. Here, we use a mouse model to isolate and sequence Dopachrome tautomerase (Dct) expressing NCCs, the precursors of melanocytes, at two key developmental stages. We classify these lineages and devise a Developmental Gene Module (DGM) scoring system to interrogate lineage state switching in melanoma samples. In bulk transcriptomes, activation of DGMs representing embryonic Schwann Cell Precursors (SCPs)—multipotent stem cells—in patient tumors predicts poor response to immune checkpoint inhibitors (ICI). Co-activation of SCP and Mesenchymal-like (Mes.) modules further correlates with resistance to MAPK inhibitors. Notably, single-cell analyses reveal that melanoma cells can simultaneously express multiple DGMs, forming “hybrid” states. Cells in a hybrid Neural/SCP state are enriched in early metastasis and ICI-resistant tumors and are insensitive to inflammatory stimuli. We demonstrate that targeting *Hdac2*, a histone deacetylase associated with this Neural/SCP hybrid state, promotes a mesenchymal-like state switch, remodels the tumor microenvironment, and sensitizes melanoma cells to TNFα and tumors to ICI therapy. Our methodology thus reveals dynamic patterns of lineage state switching correlated with melanoma tumor evolution to drive insight into new therapeutic targets.

**Teaser:** Newly identified melanoblast cell states are reawakened in metastasizing and therapy-resistant melanomas.

## Introduction

Melanoma is a cancer of the melanocyte, the pigment-producing cell in the body. Melanocytes arise from melanoblasts, precursors derived from embryonic neural crest cells (NCCs) around Embryonic (E) day 8 or from Schwann cell precursors (SCPs) from E11.5 onwards in mouse. These progenitors also give rise to neurons, glial cells, and mesenchymal derivatives; melanocytes thus share origins with diverse cell types. Melanocyte specification can be tracked through the expression of Dopachrome tautomerase (Dct), a gene encoding a pigmentation enzyme.

Melanoma cells can co-opt embryonic cell states during progression and therapy resistance[1–6], but the precise developmental states involved remain unclear. Some insights into the drivers of these NCC-like states have emerged, both intrinsic (gene regulatory networks) and extrinsic (microenvironmental cues). Rambow et al. identified RXRG as a regulator of persister cells during minimal residual disease[5], while Lu et al. demonstrated that an ALDH1A3-driven chromatin acetylation state induces NC stem cell gene expression[3], including the multipotent NCC gene, *Tfap2b*[7]. Cytokines such as IFN-γ and TNF-α also promote dedifferentiation and plasticity in melanoma cells[8–10]. Yet, the full complement of intrinsic and extrinsic regulators and how they interact remains poorly understood. The ability of melanoma cells to transition between therapy-resistant and therapy-sensitive states without new mutations mirrors the dynamic potential of NCCs[11], and lineage state switching represents a form of plasticity with major clinical consequences[12].

As in other cancers, understanding embryonic lineage specification may reveal alternative therapeutic targets [13, 14]. We reasoned that the regulatory mechanisms guiding lineage fate choices made by melanocyte precursors may be reactivated in melanoma progression (Fig. 1a). While one pioneering study deeply profiled mouse neural crest lineages across development[15], it relied on the broad Sox10:Cre-tdTomato reporter, limiting enrichment for melanocytic precursors at each stage. A second study captured human fetal and adult melanoblasts and melanocytes across many developmental stages and anatomic sites but profiled few embryonic NCC precursors of melanocytes[1]. Hence, we undertook an in-depth analysis of the melanocyte lineage at two specific stages: E11.5, with both directly developing NCCs and initial specification of SCPs and melanoblasts transmigrating across the basement membrane of the skin (mirroring vertically progressing melanoma lesions)[16, 17]; and E15.5, with melanoblasts colonizing the hair follicle niche, and SCPs establishing a stem cell niche surrounding nerves (niche establishment processes that may be relevant to metastatic colonization in cancer)[16–18]. Using a Dct-driven inducible GFP reporter mouse (iDct-GFP)[19], we enriched for melanocyte lineage cells for single-cell RNA sequencing. This enabled identification of Dct-expressing subpopulations and their linked NCC lineages, in turn leading us to devise a Developmental Gene Module (DGM) classification system to trace lineage co-option in melanoma.

**Figure 1.**
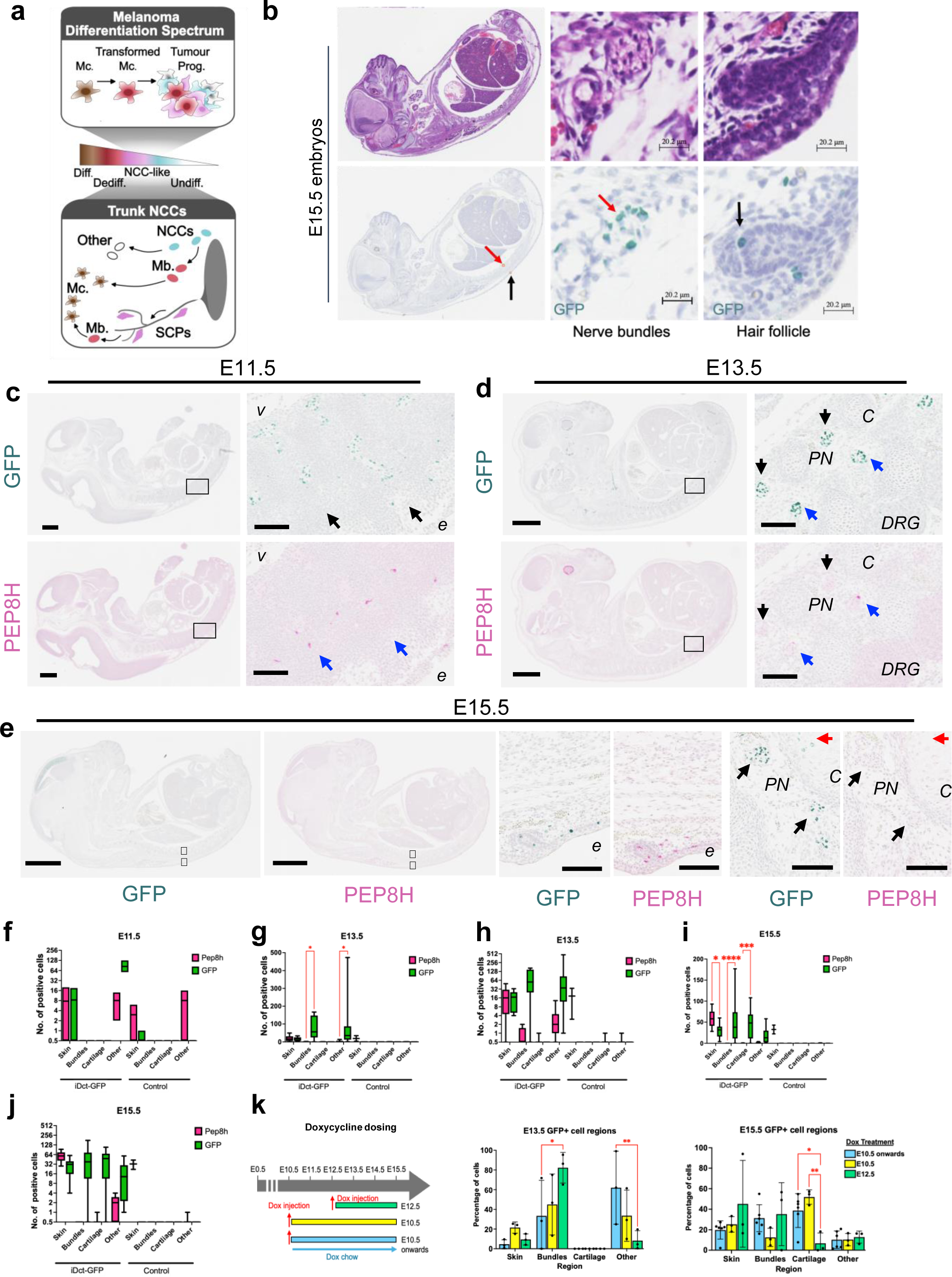
iDct-GFP reporter labels nerve-associated, migrating, and cutaneous embryonic NCCs. (**a**) Schematic of the concept of the paper. As melanomas form and progress, they dedifferentiate with cell states that represent a spectrum of dedifferentiation statuses. We hypothesize these states have parallels in normal trunk NCC development. NCCs, Neural Crest Cells; Mc., Melanocytes; Tumor Prog., tumors Progression; Diff., Differentiated; Dediff., Dedifferentiated; Undiff., Undifferentiated; Mb., Melanoblasts; SCPs, Schwann Cell Precursors. (b) Sagittal sections of E15.5 i*Dct*-GFP embryos. Top panels, Hematoxylin and eosin (H&E); Bottom panels, anti-GFP antibody (vina green). Black arrow, hair follicle; Red arrow, peripheral nerve bundle. Scale bars, 20.2μm (**c-e**), Anti-GFP (vina green) and Anti-PEP8H (magenta) staining of i*Dct*-GFP embryos. *e*, epidermis; *v*, ventral; *C*, cartilage; *PN*, peripheral nerve; *DRG*, dorsal root ganglion. (c) Black arrows, GFP+ nerve tracks; blue arrows, PEP8H+ nerve tracks. (d-e) Blue arrows, GFP+/PEP8H+ nerve bundles. Black arrows, GFP+/PEP8H-nerve bundles. (e) Red arrow, weak GFP staining in cartilage. Scale bars, E11.5, 500 μm embryo, 100 μm zoom; E13.5, 800 μm embryo, 100 μm zoom; E15.5, 2 mm embryo, 100 μm zoom. (**f-j**) Counts of GFP+ cells and PEP8H+ cells in serial sections across embryos of different stages. (**f**, **h, j**) Log scaled. (**k**) Different doxycycline dosing schedules for staged activation of the i*Dct*-GFP transgene. Red arrows, one dose intraperitoneal injection of doxycycline. Blue horizontal arrow, continuous dosing through doxycycline chow. Bar graphs of the proportion of GFP+ cells in different anatomical regions at E13.5 (left-hand graph) and E15.5 (right-hand graph). (**g**, **i, k**) Significance determined by two-way ANOVA, with Šídák’s multiple comparison test (g-i), with Tukey’s multiple comparison test (k); *, adjusted P-value < 0.05; **, adjusted P-value < 0.01; ***, adjusted P-value < 0.001; ****, adjusted P-value < 0.0001.

Surprisingly, we found that Dct+ NCCs give rise to melanocytes and glia, and a small subset of neurons and mesenchymal derivatives. Applying DGM scoring to melanoma datasets revealed that SCP-like and mesenchymal-like gene expression predicts resistance to both immune checkpoint and MAPK-targeted therapies. Moreover, we discovered that melanoma cells can exist in “hybrid” states expressing markers of multiple NCC lineages. Hybrid cells with neural and SCP features are enriched in early metastasis and Immune Checkpoint Inhibitor (ICI) resistant tumors. Finally, we identified HDAC2 as a key regulator of the non-inflammatory Neural/SCP hybrid state; its loss sensitized melanoma to TNFα and ICI, offering a potential therapeutic target in ICI-resistant tumors. In summary, by profiling embryonic melanocytic lineages at single-cell resolution and building a developmental classification framework, we identify mechanisms of melanoma plasticity and progression with direct therapeutic implications.

## Results

### iDct-GFP reporter labels cutaneous, migrating, and nerve-associated embryonic NCCs

To characterize NCCs, particularly the melanocytic lineage, we employed an iDct-GFP doxycycline-inducible transgenic mouse model that labels Dct-expressing cells. Immunohistochemistry revealed GFP+ cells in the skin and hair follicles (Fig. 1b), anticipated for the melanocyte lineage[16, 17], and surrounding peripheral nerve bundles, localized to Neural Growth Factor Receptor (Ngfr) high regions, consistent with nerve-associated Schwann cell precursors (SCPs) (Fig. 1b, Supp. Fig. 1a-c)[18].

We next compared Dct (PEP8H antibody[20]) with GFP protein expression on serial sections at embryo stages E11.5, E13.5, and E15.5. TRE-H2B-GFP single allele mice were used as controls. GFP+ and PEP8H+ cells co-localized in the skin at E11.5 and E13.5 (Fig. 1c, d, f-h). Total GFP+ cells relative to PEP8H+ cells appeared reduced at E15.5 (Fig. 1e, i-j), possibly due to differences in nuclear (GFP) compared to cytoplasmic (PEP8H) signal detection. At E11.5, GFP and PEP8H-labeled cells followed interspaced tracks consistent with dorsolateral migration (other; Fig. 1c, f), with GFP+ cells also localized more ventrally (*v*) than PEP8H+ cells, suggesting early *Dct* transcription preceding protein translation (Fig. 1c). At later stages, GFP+ cells localized near dorsal root ganglia and along nerve bundles (Fig. 1d–e), consistent with a SCP phenotype, while PEP8H+ cells could be detected in these regions, but were sparser than their GFP+ counterparts (Fig. 1d, g-j, blue arrows), indicating enhanced sensitivity of the iDct-GFP reporter. Sporadic GFP+ cells were detected between the skin and nerves, potentially representing migrating cells. GFP+ cells were also detected in cartilage at E15.5 in iDct-GFP but not control embryos (Fig. 1e, i–j), suggesting these cells may have developed from a previous Dct-expressing progenitor, underlining the importance of understanding how Dct-expressing progenitors, Dct-expressing cells, and other lineages are related.

To define lineage dynamics, we induced Dct transgene expression with either a single injected dose of doxycycline at E10.5 or E12.5 or continuous exposure (initial injection followed by doxycycline chow). Given the stability of the H2B-GFP nuclear protein, we anticipated that some signal would persist after transgene silencing. The proportion of skin GFP+ cell numbers remained consistent across induction regimens (Fig. 1k). In contrast, E12.5 induction increased the proportion of GFP+ nerve-associated cells (bundles) by E13.5, while subdermal (other) GFP+ cells decreased. These results suggest an early (E10.5) Dct-expressing progenitor origin for subdermal GFP+ populations, yet a later (E12.5) origin for nerve-associated GFP+ cells, and that either the nerve-associated cells themselves or a very recent precursor expresses Dct, in line with a potential SCP multipotent hub population. Cartilage-localized GFP+ cells were reduced following E12.5 activation, meaning they are likely to come from very early (E10.5) Dct-expressing progenitors that are more multipotent. Collectively, these findings demonstrate that the iDct-GFP model robustly labels melanoblasts and nerve-associated progenitors, and that temporal transgene activation differentially marks distinct extra-cutaneous NCC-derived populations.

### ScRNA-seq identifies distinct melanocytic and glial subpopulations in embryonic NCCs

Melanomas frequently arise in the trunk, yet previous embryonic scRNA-seq studies have largely focused on cranial and vagal NCCs[15, 21]. Using the iDct-GFP mouse model, we isolated GFP+ neural crest-derived cells at E11.5 and E15.5 from the embryo trunk (Fig. 2a; Supplementary Fig. 1d–e, Supplementary Table 1). These two stages span early NCC migration, Schwann Cell Precursor (SCP) specification, and later niche establishment in peripheral nerves and hair follicles. Cells were collected from multiple embryos per stage to mitigate litter effects and processed using the 10X Genomics platform for scRNA-seq (Supplementary Table 1). Fourteen transcriptionally distinct clusters were identified (Fig. 2b). Each embryo contributed to multiple clusters, confirmed via germline SNP-based computational genotyping (Supplementary Fig. 2a).

**Figure 2.**
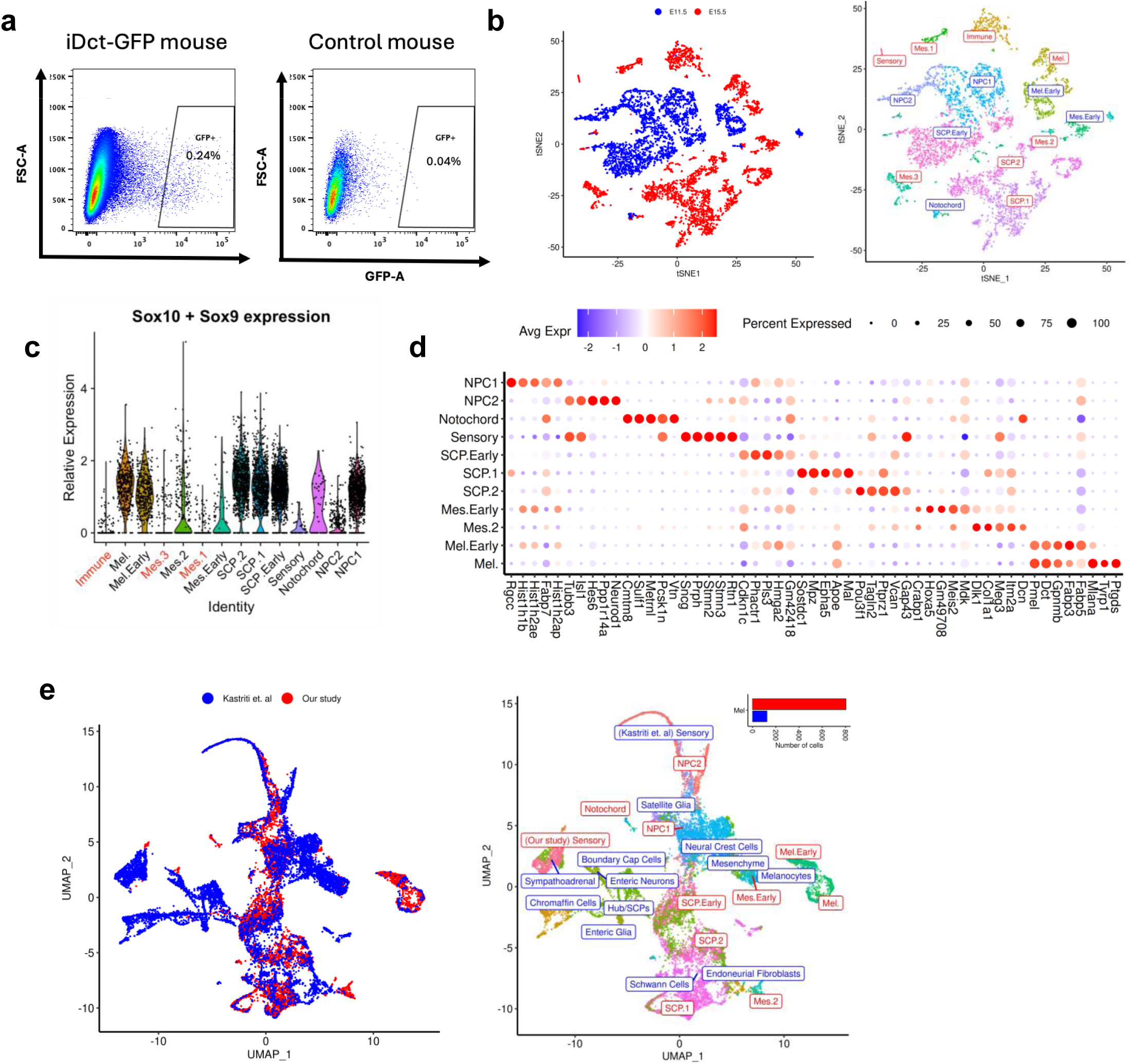
ScRNA-seq uncovers diverse subpopulations of mammalian melanoblasts with distinct phenotypes. (**a**) Fluorescence-Activated Cell Sorting of GFP+ cells from doxycycline-induced Dct-rtTA:TRE-H2B-GFP (iDct-GFP) embryos compared to TRE-H2B-GFP one allele control embryos. (**b**) tSNE embedding of GFP+ scRNA-seq data (10x Genomics). (**c**) Violin plot of cumulative Sox10 and Sox9 relative expression levels. Clusters labeled in red were removed from the study. (**d**) Dot plot of the top 5 differentially upregulated genes in each cluster. (**e**) Co-embedded UMAP plot of our data (red) and Kastriti et al. NCC data (blue). Right panel shows individual clusters. Inset shows enrichment of melanocytes in our study (red) compared to the Kastriti et al. study (blue). DGM, Developmental Gene Module; Mel., Melanocytic; Mes., Mesenchymal-like; NPC, Neural Progenitor Cell; SCP, Schwann Cell Precursor.

We enriched NCCs by selecting clusters with Sox10/Sox9 expression greater than zero and excluding non-NCC clusters, leaving eleven qualifying clusters (Fig. 2c). Marker gene analysis revealed four principal lineages: melanocytic (Mel.), SCP, neural (including sensory and neural progenitor-like cell; NPC), and mesenchymal (Mes.; Fig. 2d; Supplementary Fig. 2b; Table 1). Lineage-specific differences emerged across timepoints: E11.5 cells expressed early NCC and chromatin remodeling genes, while E15.5 cells upregulated differentiation markers (Table 1), possibly hinting that earlier NCCs likely have a greater potential, such as previously described hub cells[15].

**Table 1.**
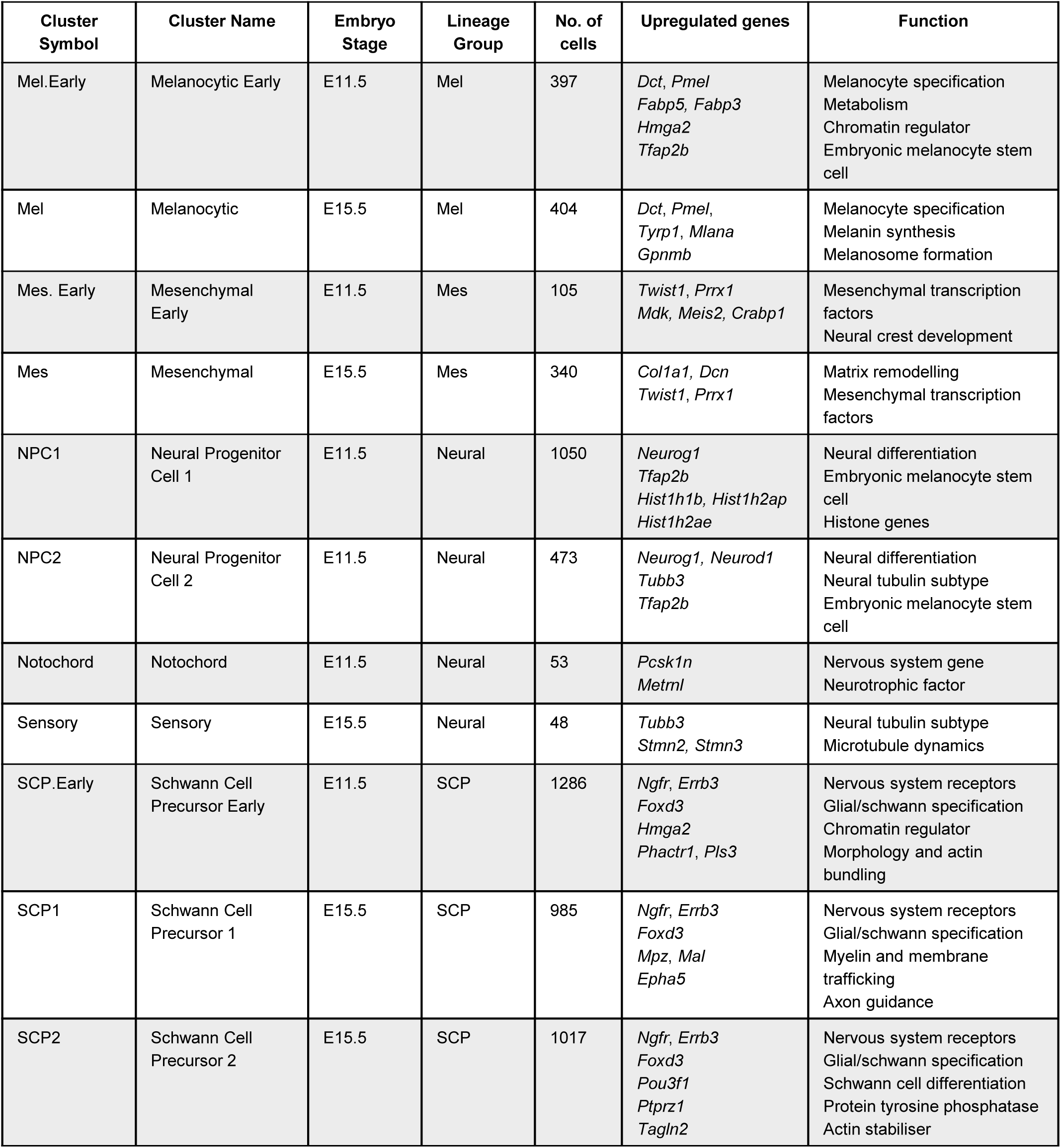

To contrast lineages enriched among GFP+ cells with those of other published neural crest populations, we integrated our dataset with Sox10-Cre-based NCC scRNA-seq (E9.5–E17.5) from Kastriti et al.[15] and tested the statistical significance of co-clustering between cells from both datasets (Fig. 2e; Supplementary Fig. 2c–d; see Methods). This should indicate which NCC lineages are closely linked to Dct-expressing progenitors, and which lineages are developmentally unrelated. Our samples were enriched for trunk NCCs and notably captured more melanocytes than Kastriti et al. Likely owing to our dissection strategy (see methods), our data lacked enteric lineages seen in Kastriti et al. The lack of sympathoadrenal lineages in our data indicated a limited overlap between these lineages and Dct-linked NCCs. Mes.Early cells co-clustered with mesenchymal populations in Kastriti et al., while Mes.2 aligned with endoneurial fibroblasts. NPC1 cells, marked by *Neurog1*, showed a higher transcriptional heterogeneity and co-clustered with neural crest cells and glial cells, indicating a possible multipotent state. Both SCP clusters at E15.5 aligned with Kastriti and colleagues’ Schwann cell clusters but remained transcriptionally distinct. The remaining neural, SCP, and melanocyte clusters in our data mapped to analogous clusters in Kastriti et al. (Fig. 2e; Supplementary Fig. 2d). In summary, the iDct-GFP model enriches for melanocytic and glial populations within NCCs, elucidating new subclusters and capturing embryonic melanoblasts at higher resolution than prior datasets, highlighting developmental plasticity between these lineages.

### Construction of Developmental Gene Modules (DGMs)

To define transcriptional programs characterizing each Dct+ neural crest state, we derived gene signatures, termed “Developmental Gene Modules” (DGM), for each of the eleven qualifying clusters (Fig. 3a). Each DGM included up to 200 genes that were (i) significantly upregulated and co-expressed within a cluster, and (ii) significantly co-expressed in the lineage (Mel., Mes., Neural, or SCP) that the cluster belonged to and was not co-expressed in other lineages and (iii) excluded cell cycle-associated genes (see Methods; Supplementary Table 3). We validated DGM specificity and sensitivity using independent scRNA-seq datasets. In Kastriti and colleagues’ Sox10+ NCC atlas[15], when we scored the cells for each DGM using AUCell, we found that the DGMs of each Dct+ lineage were upregulated in their corresponding NCC lineage and were able to correctly separate neuronal, mesenchymal, melanocytic, and Schwann/glial populations (Supplementary Fig. 2e). We further assessed the sensitivity and specificity of melanocytic DGMs using the Belote et al. dataset of fetal, neonatal, and adult human skin[1]. Both Mel. and Mel.Early DGMs were enriched in fetal and neonatal KIT+ melanocytes relative to adult cells (Supplementary Fig. 2f), consistent with their embryonic origin. These melanocytic DGMs also showed higher activity in melanocytes than non-melanocytic DGMs (Supplementary Fig. 2g), underscoring their specificity. Together, these results highlight the utility of DGMs in capturing conserved, lineage-specific gene programs across diverse scRNA-seq datasets.

**Figure 3.**
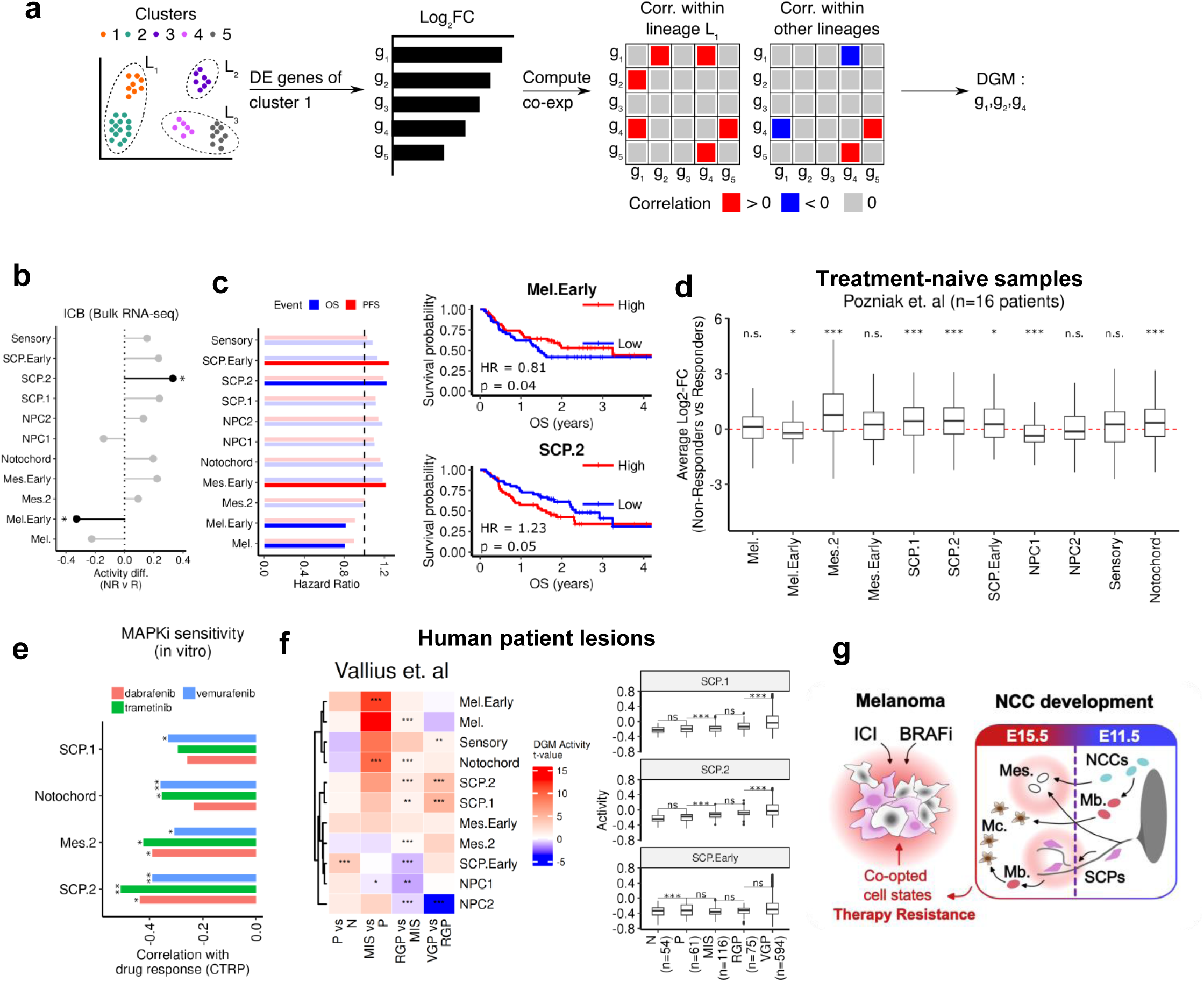
A SCP-like melanoma cell state is associated with immune exclusion in patient tumors and therapy resistance. (**a**) Procedure to derive DGMs from differentially expressed genes of Dct+ neural crest clusters belonging to different lineages. (**b**) Coefficient of linear mixed model fit (with patient cohort as random effect) of DGM activity compared to response status, where a positive value indicates higher activity in non-responders. Significance determined by t-test, * indicates p-values < 0.05. NR, Non-Responder; R, Responder. (**c**) Left: Hazard ratio of DGM activity in progression-free survival (PFS, red) and overall survival (OS, blue) of ICI therapy from Cox mixed effects regression (with cohort as a random effect) across multiple cohorts. Hazard ratios greater than 1 are positively associated with poor survival. Right: Kaplan-Meier curves of overall survival of multiple ICI cohorts[46–49], patients stratified by DGM activity. (**d**), Log2 Fold-change of DGM genes in melanoma cells before treatment from non-responders compared to responders in Pozniak et al. scRNA-seq data. Significance determined by Wilcoxon two-sided test, **** indicates p < 10^-4^, *** p < 10^-3^, ** p < 0.01, * p < 0.05 (**e**) Spearman correlation between area above drug response curve of indicated drugs across CCLE cell lines and DGM scores in bulk RNA-seq of these cell lines. Negative correlation implies that high DGM activity corresponds to lower drug-induced cell death. Significance determined by bootstrap test, * indicates p-values < 0.05. (**f**) Left: t-values of DGM activities from linear model analysis of P vs N, MIS vs P, RGP vs MIS, and VGP vs RGP stages of melanoma in GeoMx data from Vallius et al. melanoma microregions. Right: Boxplots of DGM activities across stages of melanoma progression, with the number in brackets indicating the number of microregions. Significance determined by t-test from linear mixed model with patient ID as random effect, **** indicates p < 10^-4^, *** p < 10^-3^, ** p < 0.01, * p < 0.05. N, Normal; P, Precursor; MIS, Melanoma in situ; RGP, Radial Growth Phase; VGP, Vertical Growth Phase. (**g**) Schematic to illustrate common cell states co-opted by melanoma cells and their equivalent cell state from embryonic development. Red highlighted regions depict two cell types in embryonic day (E) 15.5, Schwann Cell Precursors (SCPs) and Mesenchymal-like (Mes) that are co-opted in Immune Checkpoint Inhibitor (ICI) and targeted therapy (BRAFi) resistance. NCCs, Neural Crest Cells; Mb., Melanoblasts; Mc., Melanocytes.

### An SCP-like melanoma cell state in patient tumors is linked to therapy resistance and disease progression

Melanoma dedifferentiation is associated with poor response to immune checkpoint inhibitors (ICI)[22–24], yet the specific embryonic states recapitulated by these resistant states remain unclear. We hypothesized that melanoma cells recapitulating specific Dct-linked neural crest (NC) lineages may underlie resistance. In bulk RNA-seq from pre-treatment biopsies of anti-CTLA4 and anti-PD1-treated patients (see Methods), we found that SCP.2 was upregulated in non-responders relative to responders, while Mel.Early was enriched in responders (Fig. 3b). Consistently, SCP.2 expression predicted shorter overall survival, whereas Mel.Early was associated with longer overall survival (Fig. 3c). Focusing on progression-free survival, we found that the expression of SCP.Early and Mes.Early genes were indicative of shorter relapse times. These trends persisted in scRNA-seq data from pre-treatment biopsies of anti-PD1-treated patients, where malignant cells in non-responders showed upregulation of SCP.1, SCP.2, Mes.2, and Notochord DGMs (Fig. 3d).

Comparison of our embryonic DGMs to melanoma differentiation signatures (undifferentiated, neural crest-like, transitory, and melanocytic) defined by Tsoi et al., in their cell line panel mapped Tsoi and colleagues’ broad categories onto multiple distinct NC lineages captured in our dataset, with our DGMs offering greater lineage resolution (Supplementary Fig. 3a). As Tsoi et al. report undifferentiated melanoma cell lines being resistant to targeted therapies, we evaluated DGMs in bulk RNA-seq from untreated melanoma cell lines (CCLE) and accompanying drug response data with targeted therapies (CTRP). We found that high activity of SCP.1, SCP.2, Notochord, and Mes.2 DGMs correlated with resistance to BRAF and MEK inhibitors (Fig. 3e), while Mel. and Neural DGM activities showed no such association (Supplementary Fig. 3b). Furthermore, in BRAFi and BRAFi+MEKi-treated patients, SCP.1 activity was elevated in non-responders before therapy, especially in the combination therapy group (Supplementary Fig. 3c).

Vallius et al. have recently reported changes in melanocytic differentiation during early human melanoma progression in a large transcriptomics dataset from 62 patients[25]. The authors noted extensive heterogeneity and identified multiple histologic regions within each tumor encompassing normal (N), precursor (P), melanoma in situ (MIS), radial growth phase (RGP), and vertical growth phase (VGP) stages. Multiple microregions consisting of 100-300 cells each were sequenced from each tumor after their histologies were classified. Analyzing this data, we observed a global upregulation of DGMs in MIS microregions compared to precursor microregions and a striking stepwise increase in SCP.1 and SCP.2 activities across disease stages (Fig. 3f). Collectively, SCP and Mes. states, particularly those defined at E15.5, rather than E11.5, were associated with resistance to both immunotherapy and MAPK pathway inhibitors (Fig. 3g). These findings challenge the notion that dedifferentiation towards the earliest embryonic stage predicts resistance and suggest instead that melanoma lineage switching that co-opts characteristics of specific NC-derived lineages, especially SCPs and Mes., can drive melanoma therapy evasion.

### scRNA-seq reveals hybrid DGM cell states during early metastatic colonization

To investigate melanoma lineage plasticity during metastatic colonization, we performed single-cell RNA sequencing (scRNA-seq) on M4-BRN2 mouse melanoma cells, a highly metastatic and GFP+ subline of B2905, which were injected into the tail veins of immunocompetent mice. Analyzing lungs at 1-, 2-, and 3-weeks after injection, we observed that, as expected, tumor burden increased over time, with lesions ranging from pigmented to amelanotic, and that Sox10 expression was heterogeneous (Fig. 4a–i). We also observed metastatic nodules with a tumor-infiltrating lymphocyte component (Fig. 4c). Malignant cells were identified in scRNA-seq data by inferCNV, and non-malignant cells were annotated using Tabula Muris lung reference data (Fig. 4j, Supplementary Fig. 4a–b).

**Figure 4.**
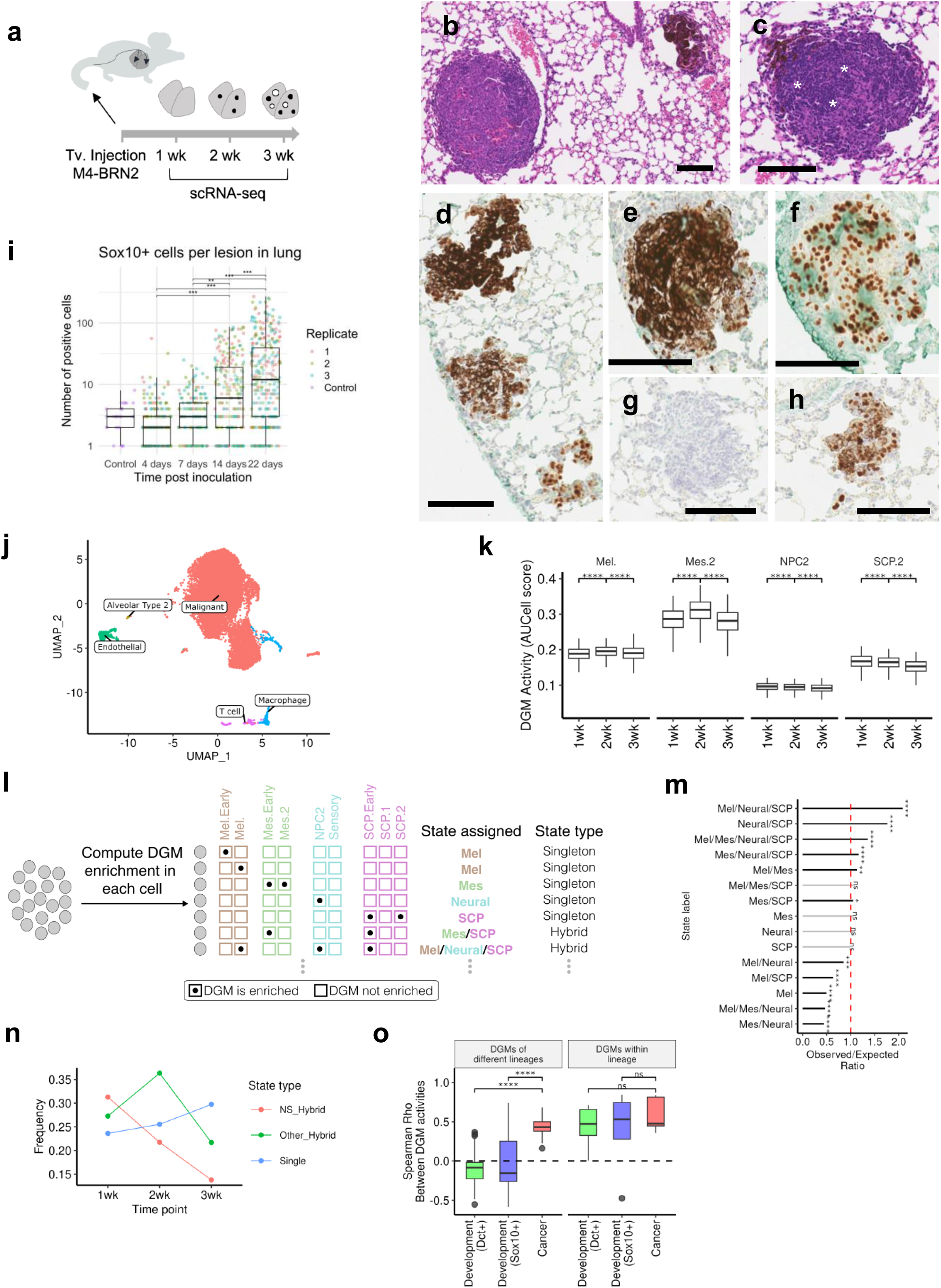
scRNA-seq uncovers dynamic DGM evolution during metastatic outgrowth. (**a**) Schematic of experimental setup. (**b-h**) Characterization of multiple pulmonary metastatic phenotypes from two-week to 22-day post-tail vein injection of M4-BRN2 mouse melanoma cells, representative examples. Bars = 100 μm. (**b-c**) Melanotic and amelanotic lesions, H&E stain. (**c**), Heterogeneous lesions with both pigmented and nonpigmented neoplastic cells and a component of tumor-infiltrating lymphocytes (*). (**d-h**) Anti-SOX10 immunohistochemistry reveals a heterogeneous pattern of nuclear immunolabeling, including some but not all lesions. (**d**) Example metastatic nodules range from intensely pigmented to heterogeneous mixtures of pigmented and nonpigmented tumor cells in some nodules (middle lesion), and relatively amelanotic nodules with distinct anti-SOX10 positive nuclei (lower nodule). (**e-f**) The same tumor nodule in serial sections is represented after anti-SOX10 without (e) and with (f) melanin bleaching to remove pigment. (**g-h**) Metastatic pulmonary nodules from the same lung section, revealing anti-SOX10 negative (f) and immunopositive (g) lesions. (**i**) Automated quantitation (QuPath) of Sox10+ lesions across timepoints. Significance determined by ANOVA, with Tukey’s multiple comparison test; *, adjusted P-value < 0.05; **, adjusted P-value < 0.01; ***, adjusted P-value < 0.001. (**j**) UMAP of all cells across all time-points obtained after batch correction and integration using Seurat. (**k**) AUCell scores of Mel., Mes.2, NPC2, and SCP.2 DGMs are all malignant cells at each time-point post inoculation. **** p < 10^-4^, *** p < 10^-3^, ** p < 0.01, * p < 0.05, two-sided Wilcoxon test. (**l**) Schematic representation of hybrid state mapping. (**m**) Ratio of observed frequency of hybrid states compared to statistical expectation by random chance in M4 cells. **** p < 10^-4^, *** p < 10^-3^, ** p < 0.01, * p < 0.05, see Methods for test details (**n**), Frequency of melanoma hybrid states at 1-, 2-, and 3-weeks post lung colonization during metastasis (**o**) Spearman’s Rho correlation score of expression of activities between DGMs of different lineages (left-hand panel), or within the same lineage (right-hand panel) across developmental datasets (Dct+, this paper) and (Sox10+, Kastriti et al., 2022) or M4 B2905 cells (this paper). **** p < 10^-4^, permutation test (**k-o**) Mel, Melanocytic; Mes, Mesenchymal-like; NPC, Neural Progenitor Cell; NT, Notochord; SCP, Schwann Cell Precursor.

We next investigated lineage changes in metastatic melanoma cells by examining the dynamic changes in DGM expression accompanying lung colonization. Analysis of lineage-specific DGM expression revealed that SCP and Neural DGMs were upregulated early but declined by week 3 (Fig. 4k, Supplementary Fig. 4c), whereas Mel. and Mes. DGMs peaked in week 2. These trends suggest that early metastatic cells transiently adopt SCP or Neural-like states, potentially facilitating adaptation. Notably, we observed cells co-expressing DGMs from multiple lineages, which we defined as a “hybrid” state (Fig. 4l), in contrast with “singleton” cells that only express DGMs from a single lineage. Such hybrid states have been reported elsewhere during tumor initiation and metastasis[26, 27]. Each hybrid state was indexed based on the lineages enriched within it. We inferred the presence of hybrid cells in our experimental metastases data and next checked if the proportion of cells in each hybrid state differed from chance (Methods). Four hybrid states co-expressing Neural and SCP DGMs (NS-Hybrids) were highly significantly overrepresented (Fig. 4m). We found that the fraction of NS-Hybrid cells significantly declined over time (Fig. 4n, Supplementary Fig. 4d-e), suggesting an adaptation role during early colonization.

Importantly, although rare (∼0.2% of cells), NS-Hybrid states were also observed in embryonic Sox10+ neural crest cells from Kastriti et al., specifically in SCPs and enteric neurons (Supplementary Fig. 4f), which aligned with the expression of multiple lineage markers in individual SCPs noted by Kastriti et al. We asked if hybrid states were more of a feature of cancer cells compared to development. We found a positive correlation between activities of DGMs within the same lineage in both cancer and development, yet only cancer cells displayed a positive correlation of DGM genes expressed between different lineages (Fig. 4o), which is consistent with hybrid states being enriched in cancer. Finally, we evaluated associations between DGM expression and plasticity in melanoma microregions profiled by Vallius et al. Using self-organizing maps, they grouped transcript signatures from tissue microregions in human primary melanomas into three “branches” and computed an entropy-based measure of plasticity from matched high-plex immunofluorescence measurement of SOX10, PRAME, NGFR, SOX9, and MART1 within each microregion. Cellular neighborhoods with melanoma cells that expressed different combinations of these lineage markers had a higher entropy. Microregions in Branch 1 had high EMT expression and higher entropy than in Branch 3. We found that SCP and Mes. lineage DGMs were significantly more active in Branch 1 microregions than in Branch 3 (Supplementary Fig. 4g), further suggesting a link between activation of these lineages and high plasticity. Together, these findings suggest that SCP-like melanoma cells co-activate other lineage programs during early metastatic and primary tumor growth, while NS-Hybrid states may underlie adaptation during early metastatic colonization.

### A non-inflammatory NS-hybrid state predicts therapy resistance and is maintained by HDAC2, which protects against TNFα-mediated apoptosis

To further assess how gene expression in hybrid and singleton lineage states evolves during metastatic colonization, we grouped the tumor cells from all three timepoints after inoculation (1, 2, and 3 weeks) based on their specific hybrid state or singleton status, identified differentially expressed genes in each group, and performed functional enrichment analysis using GSEA (Fig. 5a). Most subgroups showed a temporal shift from high interferon signaling (1 week) to EMT (2 weeks) and TNFα/NFκB activation (3 weeks), indicating a consistent inflammatory trajectory. Notably, NS-hybrid cells uniquely lacked upregulation of inflammatory pathways. This suggests that NS-hybrid cells are either refractory to inflammation or maintain a non-inflammatory phenotype. Given this, we classified hybrid states in two patient cohorts receiving anti-PD1 therapy (Pozniak et al. and Jerby-Arnon et al.[28, 29]) and found that hybrid state cells make up a significant fraction of tumor cells (Fig. 5b). More importantly, NS-hybrid cells were significantly enriched in pre-treatment biopsies of non-responders to anti-PD1 therapy (Fig. 5c), linking the NS-hybrid state to immune resistance.

**Figure 5.**
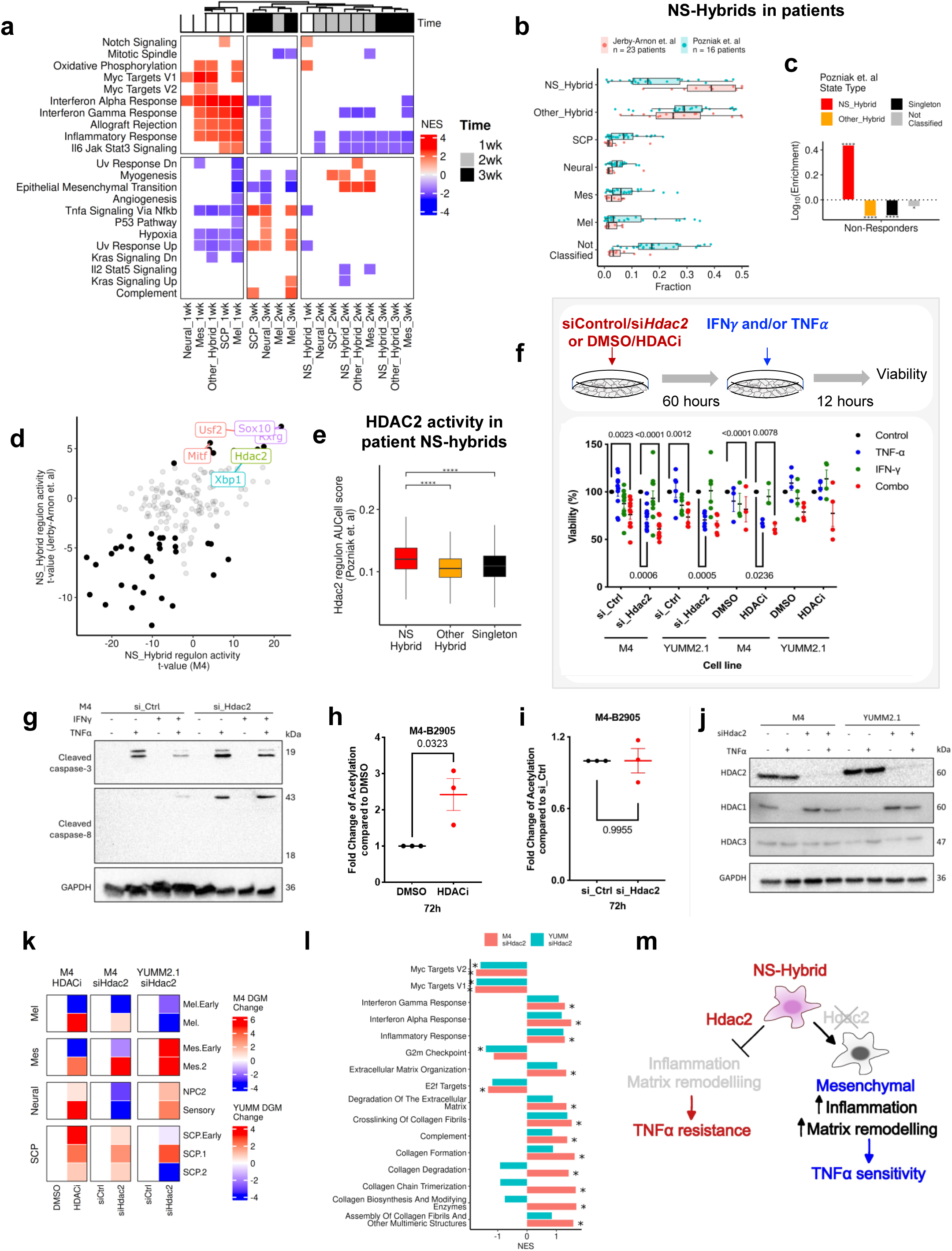
An anti-inflammatory hybrid state subtype is associated with non-response to therapy and regulated by Hdac2, which protects against TNFα-mediated cell death. (**a**) Gene Set Enrichment Analysis across hybrid and singleton states in M4-BRN2 melanoma cells during lung colonization. (**b**) Fraction of cells in each tumor from Pozniak et al., 2024, and Jerby-Arnon et al., 2018 cohorts. (**c**) Enrichment of hybrid and singleton states in treatment-naive responder and non-responder melanoma patients from Pozniak et al., 2024. **** p < 10^-4^, Fisher exact test (**d**) Regression t-values of SCENIC-inferred regulon activities in NS-hybrid cells in both mouse (M4 - this study, x-axis) and human (Jerby Arnon et al., y-axis) melanomas. Top regulons are highlighted with the colors indicating the development lineages in which regulons are specifically active (Teal: No lineage specificity, Green: Neural, Purple: SCP, Red: Mel). (**e**) HDAC2 regulon activities in treatment-naive tumor cells of the Pozniak et al. dataset. **** p < 10^-4^, two-sided Wilcoxon test (**f**) 72-hour siRNA knockdown or HDACi (0.5μm) treated cells, co-treated in the last 12 hours with cytokines, TNFα (200 ng/mL), IFNγ (1000 U/mL), or a combination of both. Percentage viability relative to siControl or DMSO control wells. Significant P-values indicated, One-way ANOVA. (**g**) Western blot of cleaved caspase-3 and cleaved caspase-8 protein in B2905 melanoma cells, treatments indicated. (**h-i**) Relative acetylation levels of HDACi-treated (h) and Hdac2 knockdown (i) mouse melanoma cells (B2905). (**j**), Western blot of HDAC1, HDAC2 and HDAC3 protein levels across mouse melanoma cell lines and treatments. (**k**) DGM activity change across HDACi and si*Hdac2*-treated mouse melanoma cells with activities in DMSO or siControl (respectively) used as a baseline for z-score calculations. (**l**) GSEA analysis of Hallmark pathways and Reactome pathways in *Hdac2* knockdown in mouse melanoma cell lines. NES: Normalized Enrichment Score. * Benjamini-Hochberg adjusted p < 0.05, permutation test (**m**) Schematic of hypotheses based on these findings. (**f-l**) si_Ctrl, non-targeting control siRNA; si_Hdac2, Hdac2 targeting siRNA; HDACi, HDAC inhibitor, entinostat.

To uncover regulators of the NS-hybrid phenotype, we applied SCENIC to i*Dct*-GFP NCC data; SCENIC infers the activity of transcriptional regulators based on the expression of their inferred target genes in each cell. Using these developmentally curated regulons, we then estimated regulator activity in cells from both human patient data (Jerby-Arnon) and our M4-BRN2 mouse metastases using AUCell. Regression analysis identified SOX10, RXRG, and HDAC2 as top regulators of the NS-hybrid state in both human and mouse melanomas (Fig. 5d). SOX10 and RXRG have previously been associated with minimal residual disease in melanoma[5], corroborating their link with a possible adaptive and therapy-resistant subtype. HDAC2 is a histone deacetylase with roles in Schwann cell lineage regulation and PD-L1 modulation[30, 31], yet further investigation of how HDAC2 may potentiate ICI therapy is warranted.

We hypothesized that HDAC2 is a candidate suppressor of inflammatory responses in NS-hybrid cells, given the lack of inflammatory gene expression enrichment in these cells. NS-hybrid cells in human patients had higher HDAC2 activity compared to other hybrid cells and singleton cells (Fig. 5e), corroborating HDAC2 as a key gene in NS-hybrid cell function. Using the M4-BRN2 parental cell line, B2905, and an additional mouse melanoma cell line, Yumm2.1, that each had different SOX10 expression statuses (Supplementary Fig. 5a-b), we performed *Hdac2* knockdown (siRNA), followed by TNFα stimulation. We confirmed TNFα action through immunoblotting of phosphorylated p65 (a key part of the NF-κB pathway and activated by TNFα; Supplementary Fig. 5a-b). This led to ∼30% viability loss compared to non-stimulated cells, where non-targeting control siRNA-transfected cells (siControl) stimulated with TNFα demonstrated no change in viability compared to non-stimulated cells. IFNγ stimulation alone had no effect across modalities (Fig. 5f). This TNFα-specific cytotoxicity correlated with increased cleaved caspase-3 and -8, implicating apoptosis (Fig. 5g).

Next, we asked if we could mimic this *Hdac2-*KD-driven TNFα-sensitivity phenotype using a small-molecule inhibitor approach, as this may be directly translatable to the clinic. Entinostat is a Class I pan-HDAC inhibitor (HDACi) inhibiting HDACs 1, 2, and 3. Entinostat has been tested in phase I and II clinical trials against advanced malignancies and in combination with immune checkpoint inhibitor therapies[32, 33]. Like *Hdac2* KD, B2905 cells treated with HDACi exhibited a TNFα-specific loss of viability, which was not exacerbated with combination treatment (Fig. 5f). However, in contrast, HDACi-treated Yumm2.1 cells tolerated TNFα, suggesting that *Hdac2* loss alone may be a more efficacious target across multiple cell lines than inhibition of all 3 HDACs simultaneously. Chromatin acetylation was increased with HDACi treatment, as would be expected following inhibition of HDACs 1-3 (Fig. 5h), validating the drug’s activity. Yet chromatin acetylation levels remain unchanged in *Hdac2* KD cells (Fig. 5i), which suggests other histone deacetylase proteins may be upregulated to compensate for the loss of *Hdac2*. Western blot analysis revealed that HDAC1 was upregulated following *Hdac2* KD (Fig. 5j), suggesting a mechanism through which global acetylation levels remain unchanged. However, regardless of acetylation levels, HDAC2 loss was sufficient to sensitize cells to TNFα, suggesting HDAC2 may have different targets than HDACs 1 or 3.

RNA-seq of *Hdac2* KD or HDACi-treated cells revealed widespread DGM alterations, especially a strong upregulation of the mesenchymal Mes.2 module (Fig. 5k) and indicated a shift toward a mesenchymal state upon HDAC2 depletion. Universally, TNFα treatment led to a dramatic downregulation of all SCP gene modules (Supplementary Fig. 5c), in line with negative correlations between SCP expression and TNFα pathway activation in lung metastases. Functional enrichment analysis using GSEA revealed activation of interferon response and inflammatory pathways in *Hdac2* KD cells, consistent with a role for HDAC2 in mediating a non-inflammatory phenotype (Fig. 5l). We also found an upregulation of matrix remodeling pathways in *Hdac2* KD cells. Thus, while it is uncertain if HDAC2 drives the NS-hybrid state per se, it may repress both a transition to a mesenchymal state and sensitivity to inflammatory stimuli. Loss of HDAC2 disrupts this balance, promoting mesenchymal differentiation, immune pathway activation, and vulnerability to inflammatory cues (Fig. 5m).

### HDAC2 Loss reprograms the melanoma microenvironment to re-sensitize tumors to anti-PD-1 therapy

HDACi has shown potential in the clinic in combination with PD-1 blockade therapy, though results have been mixed[34]. Based on the *in vitro* effects of *Hdac2* KD above, we hypothesized that HDAC2 loss would elicit lineage changes in tumor cells, impacting their interaction with the stromal microenvironment, potentially sensitizing tumors to anti-PD-1 therapy. To test this, we transplanted *Hdac2* KD melanoma cells subcutaneously in mice. siRNA knockdown allows a fast and reversible loss of *Hdac2* gene expression (up to 10-14 days), exposing immediate tumor cell responses to HDAC2 loss without the confounding effects of long-term cellular adaptation mechanisms. Moreover, with this approach, we could interrogate how loss of HDAC2 specifically impacts tumor colonization at a primary site (under the skin). Subcutaneous tumor growth assays also allow for accurate pre-clinical testing of tumor growth dynamics and therapy response.

Following tumor implantation (either siControl or si*Hdac2*), immunocompetent mice were treated with either IgG control or anti-PD-1 (Fig. 6a). Both control and *Hdac2* KD tumors grew equally under IgG control treatment. After anti-PD-1 treatment, both control and *Hdac2* KD tumors regressed; however, the *Hdac2* KD tumors showed significantly greater regression, remaining complete for up to 2 weeks after stopping treatment, before relapsing. In contrast, control tumors treated with anti-PD-1 exhibited only partial regression and relapsed soon after treatment stopped (Fig. 6b and Supplementary Fig. 5e). We consider Hdac2 KD IgG-treated tumors as treatment-naive tumors likely to respond to ICI therapy, whereas siControl IgG-treated tumors represent treatment-naive partial responders (based on Fig. 6b). Both anti-PD-1 tumors are considered relapsed tumors. In summary, transient loss of HDAC2 at tumor cell transplantation sensitized tumors to anti-PD-1 therapy.

**Figure 6.**
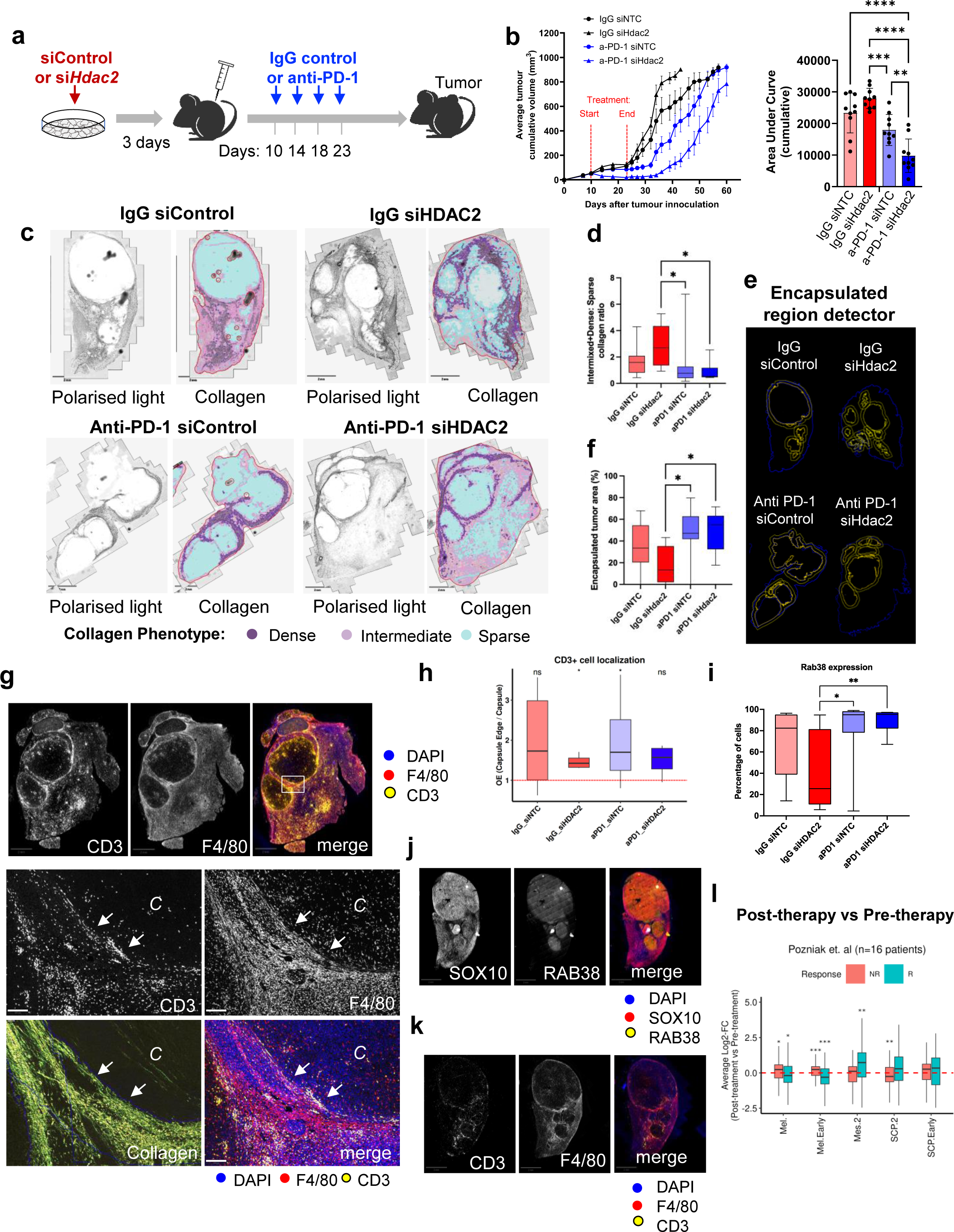
HDAC2 Loss reprograms the melanoma microenvironment to re-sensitize tumors to anti-PD-1 therapy. (**a**) Schematic of treatment schedule. (**b**) Left: Tumor growth curves, treatment indicated. Right: cumulative Area Under Curve extrapolated from tumor growth curves. (**c**) Left-hand panels, Optical density sum projected image of picrosirius red stained tumor sections visualized under polarized light. Right-hand panels, Qupath AI-generated detection of sparse, intermediate, and medium collagen densities based on the optical density sum images. (**d**) Ratio of tumor area comprised of dense/intermediate collagen versus sparse collagen. (**e**) Example of regions identified by the automated capsule analysis algorithm (Qupath). (**f**) Proportion of tumor area encapsulated. (**g**) Tissue sections stained with anti-CD3 and anti-F4/80 alongside a picrosirius red stain of a serial section visualized in polarized light (Collagen). *C*, Capsule; arrows, edge regions of high CD3+ cells, high F4/80+ cells alongside collagen fibrils. (**h**) Fold change of proportion of tumor CD3+ cells at the capsule edge/ in the capsule. (**i**) Percentage of Rab38+ cells per tumor per treatment. (**j**) Immunostaining of tumors with anti-RAB38 and anti-SOX10 antibodies. (**k**) Immunostaining of tumors with anti-F4/80 and anti-CD3 antibodies. **(l)** Log2 Fold-change of DGM genes in melanoma cells of post-treatment cells compared to pre-treatment cells amongst responders and non-responders in Pozniak et al. scRNA-seq data. (**d, f**) Kruskal Wallis test, (**h**) two-sided Wilcoxon test, (**b, i**) One-way ANOVA, Tukey’s multiple comparison test; *, p< 0.05; ** p < 0.01, *** p<0.001, ****p<0.0001. (**l**) two-sided Wilcoxon test; **** p < 10^-4^, *** p < 10^-3^, ** p < 0.01, * p < 0.05. (**f-l**) M4, B2905 mouse melanoma cells.

We reasoned that transient HDAC2 loss might lead to lasting changes in tumor evolution, making tumors more sensitive to immune clearance triggered by ICI therapy. We then examined how transient HDAC2 loss affected tumor cell phenotypes and the tumor microenvironment at the time of transplantation. Using multiplexed tissue profiling, we analyzed tumor lineage phenotypes and tissue characteristics. Based on our RNA-seq data (Fig. 5k-l), which showed increased activation of the mesenchymal module and significant changes in collagen biology after HDAC2 depletion, we investigated whether there were observable differences in tumor collagen content—a key feature of the tumor microenvironment—between siControl and siHdac2 knockdown tumors treated with either IgG control or anti-PD-1. Picrosirius red staining revealed different collagen phenotypes, with regions of dense, intermediate, and sparse collagen. The ratio of dense/intermediate collagen to sparse collagen was higher in IgG-treated *Hdac2* KD tumors (which were treatment-sensitive) compared to both anti-PD-1-treated groups (which relapsed). Conversely, siControl tumors (partial responders) showed no significant difference across any treatment groups (Fig. 6c-d, Supplementary Fig. 6). Many regions of sparse collagen were surrounded by borders of denser collagen; these features were regarded as encapsulated regions. The encapsulated tumor area was smaller in treatment-naive siHDAC2 tumors compared to siHdac2 relapsed tumors, while no difference was observed in siControl tumors before and after treatment (Fig. 6e-f). This encapsulated phenotype could exclude entry of immune infiltrates and thereby help convey resistance to ICIs[35]. Therefore, we hypothesize that the presence of these encapsulated regions may alter the organization of tumor-infiltrating lymphocytes (TILs) within the tumor and ultimately impact resistance to ICI therapies. We observed that CD3+ T cells and macrophages colocalized at the circumference of these collagen capsules (Fig. 6g-h, Supplementary Fig. 6). Thus, the presence of capsules results in regions of lower T cell infiltration, mimicking an immune cell exclusion phenotype.

We next asked how expression of lineage markers altered across tumors. While SOX10 expression was unaltered, we observed that tumors with greater numbers of capsules, such as following anti-PD-1 therapy, had significantly enriched RAB38 expression (a melanocytic marker upregulated in SCP.1 cells) compared to tumors with fewer capsules (*Hdac2* KD, IgG treated). This suggests relapsed tumors were more melanocytic post anti-PD-1 therapy, but less melanocytic in *Hdac2* KD, IgG-treated modalities (Fig. 6i-j; Supplementary Fig. 6; Supplementary Fig. 7a). This result initially surprised us as high Mel.Early expression before anti-PD1 therapy was associated with response and better survival (Fig. 3b, c). We, therefore, asked if, like our mouse data suggests, anti-PD1 treatment increased melanocytic gene expression in relapsed tumors[29]. We computed log-fold changes of genes in DGMs in post-treatment samples of non-responders and responders compared to their pre-treatment baselines and found that non-responders indeed up-regulated melanocytic DGMs post-therapy, whereas responders down-regulated melanocytic modules (Fig. 6l; Supplementary Fig. 7b). This parallels prior work reporting an unexpected dedifferentiation phenotype in melanoma cells following therapy in non-responding patients[10]￼. Altogether, our work demonstrates the dynamic nature of lineage state switching and its relationship to the tumor microenvironment, including collagen remodeling and cytokine stimulation. SCP and Mes. lineage DGMs were activated in early melanoma microregions with immune infiltration as well as in TCGA samples infiltrated by CD8 T cells (Supplementary Fig. 7c-d). This link between lineage switch and inflammation may be a fundamental feature of the Dct+ neural crest lineage, as Mes.2 and SCP.1 NCCs had higher inflammatory signaling compared to other embryonic cell clusters in our data (Supplementary Fig. 7e). Also, analysis of gene expression data from mouse melanocytes irradiated by [19]￼, revealed activation of SCP.2 module expression at a timepoint post UVB treatment (Supplementary Fig. 7f) where upregulation of immuno-evasion genes had been previously reported.

## Discussion

Our study refines longstanding observations that cancers, including melanoma, reactivate developmental programs. By deeply profiling Dct-expressing cells at two key stages in mouse embryogenesis, we identified specific neural crest cell (NCC) states, particularly late embryonic (E15.5) Schwann cell progenitors (SCPs), that are re-acquired during melanoma progression. These SCP-like states are clinically relevant: they correlate with immune checkpoint inhibitor (ICI) resistance independently of traditional markers like tumor mutation burden[36] or tumor microenvironment profiles (TME)[37], and are enriched in invasive primary tumors and early metastases. Moreover, because the SCP signature reflects a natural embryonic lineage, it offers a biologically grounded framework to explore the signaling, niches, and plastic potential associated with these states in melanoma.

As plastic adaptation is a dynamic process that has been traditionally difficult to predict, let alone manipulate, we propose that comprehensive longitudinal mapping of melanoma lineage states enabled by our iDct-GFP embryonic dataset may help bridge this gap in precision oncology. Our dataset reveals greater diversity in the Dct+ neural crest lineage than previously recognized, connecting early progenitors to both glial and melanocytic fates predominantly. This resource helps define coordinated gene programs (DGMs) in melanoma with greater resolution than traditional approaches based on a small number of lineage markers. Notably, our lineage signatures extend the developmental subtyping frameworks of Tsoi et al.[6] and Belote et al.[1] by identifying precise Dct-linked embryonic lineages reactivated in melanomas. While Belote et al. largely profiled melanocytes from embryonic and adult stages, we did not sequence Dct+ lineages from adult mice, a limitation that, when addressed, could help span the full differentiation trajectory represented in our DGMs.

An open question in melanoma biology is how the interplay between extrinsic and intrinsic determinants drives melanoma evolution. Our metastatic colonization data reveal a temporal switch in inflammatory signaling within tumor cells: an early phase marked by interferon activation (including IFN-γ) and SCP-like gene expression, followed by a transition toward TNFα/NFκΒ signaling. IFN-γ plays a dual role—initially promoting immune targeting but eventually enabling immune evasion in tumor cells[38, 39]. This lineage-intrinsic response is mirrored in melanocytes, where UVB exposure induces both IFN-γ signaling[19] and SCP gene expression, perhaps linking an early stress response in melanocytes to the SCP module. Notably, SCP DGM activity is upregulated in metastatic cells at 1-week post-injection compared to later timepoints, preceding the inflammatory switch. We also identify two populations of SCPs at E15.5, of which SCP.1 cluster cells have enhanced inflammatory pathway activation and SCP.2 cluster cells have no inflammatory pathway activation. Thus, inflammation could play an important role in distinguishing biologically relevant SCP subtypes. We propose that SCP-like sub-states may mediate IFN-γ signaling, contribute to immune refractoriness, and enable phenotypic plasticity in melanoma. What adaptive advantage expression of SCP genes affords the tumor cells is yet to be determined, but potentially, expression of SCP genes could enable tumor cells to suppress antigen expression and/or presentation or enable greater plastic potential to permit a switch to immune-refractory states. For example, IFN-γ signaling has been shown to upregulate Nerve Growth Factor Receptor (NGFR) activity in cells. NGFR is a marker of SCPs, and NGFR signaling has been linked to immune therapy refractory, stem-like melanomas[40]. Taken together, an IFN-γ-SCP axis may signify a potential melanoblast lineage-induced mechanism of immune cell evasion.

At single-cell resolution, we found that a subset of melanoma cells co-express markers from multiple Dct-linked lineages simultaneously, forming hybrid states. These are rare in normal development but do occur, especially among SCPs and enteric neurons. We propose that, as in development[15, 41], hybrid states in melanoma reflect lineage plasticity. One particularly notable state—NS-Hybrid—is enriched in SCPs and neural lineages and is prevalent in early metastasis and ICI-resistant tumors. While the mechanisms driving these hybrid phenotypes remain unclear, the role of SCPs as multipotent progenitors, their increased activity during progression, and their enrichment in regions of high spatial entropy all suggest a functional role for the SCP state in mediating this hybridity. SCP module activation may define a permissive, hub-like state facilitating cell-state transitions, similar to common cell fate decision routes hypothesized in dynamical theories of cell decision-making[42], and aligning with recent descriptions of SCP hub states[15] and cyclical fate restriction models in zebrafish[41].

We highlight active NS-Hybrid regulons in both mouse and human melanomas as promising therapeutic targets. In our model, high HDAC2 activity was linked to the non-inflammatory phenotype of NS hybrid cells during lung colonization. Transient knockdown of *Hdac2* expression increased melanoma cell sensitivity to TNFα-induced apoptosis, possibly by reduced expression of C-FLIP, a known anti-apoptotic factor previously implicated in pancreatic cancer[43]; alternatively, it may be due to a natural sensitivity to TNFα of a mesenchymal-like state. HDAC2 also bestowed resistance to the tumor growth suppressive action of ICI. This is likely multifaceted, reflecting changes in sensitivity to TNFα released as tumors become inflamed, but also in part due to HDAC2-mediated deacetylation of PD-L1, affecting its nuclear translocation[30]. Moreover, in pancreatic cancer, HDAC2 has been linked to metastasis[44].

Moreover, tumors evolving from HDAC2-depleted tumor cells exhibited reduced encapsulation by cross-linked collagen and a concomitant reduction in RAB38 expression (a melanosome biogenesis and trafficking protein also implicated in metastasis)[45]. Collagen encapsulation of tumors can alter T cell distribution[35], excluding T cells from entering the encapsulated masses, rendering the tumor more resistant to ICI. Accordingly, the less encapsulated HDAC2-depleted tumors are more sensitive to ICI. While collagen capsules are likely assembled by tumor-resident fibroblasts, our RNAseq data suggest that tumor cells themselves may contribute. The relative importance of stromal to tumor-derived collagen and its regulation remains unclear; interactions among tumor cells, fibroblasts, T cells, and macrophages likely shape this architecture. Future studies should explore how HDAC2-depleted cells modulate tumor evolution to reduce encapsulation and whether this can be therapeutically exploited.

Understanding the precise chromatin acetylation program that is executed in HDAC2-high or HDAC2-low contexts and how these differ from HDAC1 or HDAC3 is crucial, particularly considering HDAC inhibitors under clinical evaluation. The higher sensitivity of HDAC2 to TNFα relative to its family members, which our results suggest, could guide future drug development. More broadly, shifts in chromatin acetylation are associated with stemness and therapy resistance. Particularly, ALDH1A3-driven metabolic acetylation of histone H3 has been linked to upregulation of the embryonic gene[3], *Tfap2b*[7], critical to drug-refractory melanoma.

Finally, DGM-based lineage mapping has shed light on how melanoma lineage states evolve in response to therapy. Melanocytic, differentiated melanomas in a pre-treatment setting are generally more sensitive to ICI; nonetheless, relapsed tumors adopt a more melanocytic phenotype post-treatment. One possible resolution to this paradox is that a melanocytic identity confers a selective advantage only after the tumor has reconditioned the TME to evade immune attack. For instance, we observed that relapsed tumors in our model exhibited increased RAB38 expression and were encased in dense collagen capsules that excluded T cells from the tumor mass. We speculate that the tumor must first remodel its microenvironment—potentially by forming a collagen capsule—to physically shield these otherwise immunogenic melanocytic cells from immune surveillance. In this scenario, once immune pressure is mitigated by encapsulation and other TME changes, reactivating a melanocytic program becomes a selective advantage post-therapy.

In summary, through analysis of curated single-cell developmental and metastasis datasets and comprehensive analyses of developmental lineages in published data, we reveal how inflammatory signals shape developmental plasticity in melanoma. We have generated rich datasets and developed signatures to fully map the contribution of fundamental, conserved embryonic neural crest lineage pathways to melanoma progression. We explore, for the first time, how single melanoma cells can simultaneously co-opt multiple neural crest pathways and how these hybrid states contribute to tumor evolution. These data, and the tools we provide, help us gain new insight into mechanisms of melanoma plasticity, metastasis, and therapeutic resistance.

## Materials and Methods

### Mouse models and melanoma cell lines

Melanoblasts were isolated from the Dct-rtTA; tre-H2B-GFP (*i*Dct-GFP; Frederick National Laboratory for Cancer Research) mouse model. Embryonic development was timed based on the number of days post coitum. Pregnant females (Fig. 1b, 2, and Supplementary Fig. 1) were placed on a doxycycline-enriched diet (200mg/Kg) to activate expression of GFP. For lineage tracing experiments, either doxycycline chow or intraperitoneal injection of doxycycline (80µg per gram body weight) was administered by the dosing schedule indicated (Fig. 1k). Melanomas were derived from the following mouse melanoma models: M4, C57BL/6 male background, with Hgf-tg transgenic allele. UV was used as the tumor-inducing carcinogen; M4 mice were treated at postnatal day 3[46]. Cell lines derived from this model (B2905) were transduced with Pol2-ffLuc-IRES-H2B-eGFP, then cycled through mice via intracardiac injection and isolated from the brain and cultured[47]. The M4-BRN2 cell line derivative has been cycled through mice twice to enrich cells of high metastatic potential. Experimental metastasis studies were performed using a filtered, single-cell suspension in phosphate-buffered saline (PBS). 1×10^6 M4-BRN2 cells were injected in a 100-µl volume into the tail vein of 6–8-week-old Glowing-Head mice (Frederick National Laboratory for Cancer Research) x C57BL/6N mice (Charles River, Frederick National Laboratory for Cancer Research), making an F1 cross. Lungs were perfused with PBS through the circulation before removal for sample preparation. Lungs were removed from mice at the timepoints indicated. B2905 and YUMM2.1 cells (provided by Richard Marais, The University of Manchester) were maintained in RPMI-1640 (Sigma– Aldrich) supplemented with 10 %v/v fetal bovine serum (FBS; Life Technologies) and 1% l-glutamine (Gibco). All cells were maintained under standard conditions at 37 °C in a 5 %v/v CO_2_ humidified incubator and passaged before reaching confluency. Cell lines were authenticated by STR profiling, and cultures were routinely tested for mycoplasma contamination by PCR and deemed to be uninfected.

### Single Cell – Sample Preparation, Sorting and Oligo-Labeled Antibody Staining

For melanoblast preparation, multiple litters were used per time point (3 litters at E11.5 and 4 litters at E15.5), and one embryo per litter was used for melanoblast isolation. Trunk melanoblasts were isolated from a preparation that excluded the head, arms, legs, and internal organs, but retained the dermis, epidermis, spinal cord, and associated nerve and muscle tissue. Enzymatic dissociation of melanoblasts was as follows: tissue was cut finely and digested with an incubation of 30 minutes at 37 °C with a digestion medium of RPMI media and 200 U/mL Liberase TL (Roche). Following digestion tissue was kept cold on ice. Physical dissociation of melanoblasts was performed using 50 μm green medicons (BD Bioscience), E11.5 and E15.5 embryos processed on the medimachine for 1.5 mins and 2 mins, respectively. Cells resuspended in a medium/10% fetal bovine serum/DΝase I and spun at 300xg for 5 mins. Cells were washed in a solution of PBSA (PBS and 1% Bovine Serum albumin) and filtered through a 30μm filter for cell sorting. Cells were incubated for 30 minutes with LIVE/DEAD Violet stain (ThermoFisher) as per the manufacturer’s instructions or were stained with DAPI immediately before sorting.

For Lung metastasis preparation, 3-4 lungs were used per time point (1-, 2-, 3-weeks post-injection). Each lung was perfused with PBS via the circulation and cut into small pieces. A solution of 0.5 μg/μL Liberase TL (Roche) in media (RPMI/DMEM) was added to the tissue pieces in a Miltenyi C tube. The chunks were processed on the Miltenyi tumor dissociator, then incubated at 37°C for 30-60 min. The chunks were processed on the tumor dissociator once more and then transferred to a solution of RPMI containing 20% FBS and 0.5-mg/ml DNase (Sigma). Cells were filtered using a 70μm filter and the pellet resuspended in ACK lysis buffer for 5-10 minutes. Cells were washed, filtered, and stained with LIVE/DEAD Violet stain (ThermoFisher) as per the manufacturer’s instructions, or were stained with DAPI immediately before sorting.

Oligo-labeled antibody staining was done on both melanoblasts and melanoma cells using the TotalSeq™-A Antibodies and Cell Hashing with 10x Single Cell 3’ Reagent Kit (BioLegend). The HashTag Oligonucleotides used for the analysis are mHTO01, mHTO02, mHTO04, and mHTO05. TotalSeqA from BioLegend Staining was performed as per the manufacturer’s guidelines using TruStain FcX™ PLUS (anti-mouse CD16/32) (BioLegend) as a blocking agent. For cell hashing antibodies, we used 0.5 μg per 1 million cells, followed by 3 washes before 10x Genomics sequencing.

### Fluorescence-activated cell sorting (FACS)

iDct-GFP-positive melanoblasts/melanocytes and eGFP-positive melanoma cells were sorted on a BD FACSAria IIu (BD Biosciences) cell sorter. Cells were initially identified on forward scatter (FSC) and side scatter (SSC). Cell doublets and aggregates were excluded using light scatter pulse width. Cells were sorted based on GFP/eGFP expression and viability as determined by staining with DAPI or LIVE/DEAD Violet stain (ThermoFisher). Embryos of the same developmental age that were heterozygous for the TRE-H2B-GFP gene but lacked the Dct-rtTA gene or derived from the FVB were used as negative controls for melanoblasts. Lungs from C57BL6 wild-type strain mice that were uninjected were used as negative controls for metastatic studies. Due to low sample cell number, reanalysis of sorted cells was not usually done, but representative post-sort analyses confirmed that initial frequencies of 0.6–4.4% GFP-positive cells were enriched up to 97% following the sort.

### Single Cell – Partitioning and Library Preparation

Prepared antibody-label multiplexed single cell suspensions were washed with cold PBS with 0.04% BSA by centrifugation at 300g and gentle resuspension in fresh buffer. Cell counts and viability measurement were performed on each cell suspension using an automated cell counter with propidium iodide and acridine orange dyes (LunaFL, Logos Biosystem). Cell concentrations were adjusted for loading onto the 10x Genomics Chromium platform using the 3’ v3 gene expression chemistry (10x Genomics), targeting approximately 6,000 cells per sample. Single-cell RNA-Seq libraries were prepared according to vendor recommendations.

### Single Cell – Sequencing

Sequencing was performed on the Illumina NextSeq 550 (for embryonic melanoblasts samples) or NextSeq 2000 (for melanoma metastasis samples), with separate sequencing runs for gene expression libraries and associated antibody-oligo libraries. Single-cell gene expression libraries were sequenced with a 28bp read to identify cell barcodes and unique molecular indices, an 8bp read for sample indices, and a 98bp read to identify cDNA insert. For antibody-oligo libraries, a shorter insert read was used. Samples were multiplexed for sequencing based on experimental groups, and reads were combined from multiple sequencing runs to achieve a target greater than 50,000 reads per cell on average for gene expression and at least 10,000 reads per cell for antibody-oligo libraries for all samples. >4000 melanoblasts were isolated collectively from the multiple embryo donors at each stage. Sequencing depth was excellent, >71,000 mean reads per cell, yielding >12,000 median Unique Molecular Identifiers (UMI) per cell and identifying > 3,700 median genes per cell.

### Single Cell – Data Processing

Data was processed using the 10x Genomics CellRanger pipeline to demultiplex reads and then align reads to a GRCh38 reference for scRNA-Seq data (cellranger v6.1.0 against reference refdata-cellranger-mm10-3.0.0 (for embryonic melanoblasts samples) augmented with the GFP sequence and cellranger v6.1.01 against refdata-gex-mm10-2020-A (for melanoma metastasis samples) augmented with the GFP sequence. UMI-adjusted aligned reads were used to generate a single-cell barcode and gene expression matrix, and an antibody-oligo count matrix that was used for downstream analysis.

### Histology and Immunolabelling

All embryo histology was undertaken by Histoserv, Inc. (Germantown, USA). Formal-Fixed Paraffin-Embedded (FFPE) slides were deparaffinized in graded alcohols and brought to water. For slides that required bleaching, bleaching was performed in a heated bath with 3 %v/v H_2_O_2_ in PBS for 30 minutes at 60 °C. Antigens were then retrieved at 100 °C for 20 minutes. After retrieval, slides were blocked with a peroxidase block, incubated with the primary antibody, and detected with an HRP-conjugated goat-anti-rabbit secondary. HRP was then visualized with either DAB, magenta, or Vina Green chromogens. For multiplex slides, the staining was repeated on the same slides, beginning with another round of retrieval, following the same detection protocol, and completed by visualizing the staining with an alternative chromogen. Upon staining completion, slides were dehydrated and cleared in rapid changes of graded alcohols and xylene (or clearite for Vina Green) and mounted with permanent mounting medium. Primary antibodies, concentrations used, and hosts: Sox10 - Cellmarque #383R, 1:300; Rabbit host; Col VI - Novus #NB120-6588, 12ug/ml; Rabbit host; GFP - Genetex #20290, 1:400; Rabbit host; Zeb1 - Cell Signaling #70512, 1:100; Rabbit host. For lineage tracing embryo section staining: PEP8H antibody was gifted by Vincent Hearing[20] using pH9 retrieval, 1:5000 dilution, magenta chromagen. GFP antibody, Abcam ab183734, pH6 retrieval, 1:3200 dilution (0.043ug/ml), Vina green chromagen.

Tumors were formalin-fixed (4% formaldehyde in PBS) for a minimum of 24 hours at 4 °C, then kept in 70 %v/v ethanol at room temperature until paraffin embedding (FFPE). FFPE tumors were sectioned at 5 µm and randomised on frosted slides.

### Mouse tumor implant study design

Mice were housed in the Biological Services Facility of The University of Manchester on a 12/12 h light/dark cycle, and given unlimited access to food (Bekay, B&K Universal, Hull, UK) and water. All procedures were approved by the University of Manchester’s animal welfare ethical review board and performed under relevant Home Office licences according to the UK Animals (Scientific Procedures) Act, 1986. Female, 8–12-week-old C57BL/6 mice were purchased from ENVIGO and allowed at least 1 week to acclimatize. M4-B2905 cells (2 × 10^6^ cells) in 100 µL serum-free RPMI-1640 were subcutaneously injected into the left flank of mice under isoflurane anaesthesia. Tumor size (calculated by multiplication of height, width, and length caliper measurements) and mouse weight were monitored three times per week (every 2–3 days). When tumors reached an average volume of 80–100 mm^3^, mice were administered up to four doses of 300 µg of α-PD-1 antibody (BioXCell) or rat isotype control antibody IgG2a (BioXCell) in 100 µL InVivoPure pH 7.0 Dilution Buffer (BioXCell) via intraperitoneal (i.p.) injection administered at 3–4-day intervals. Mice were culled once tumors reached 800 mm^3^, our pre-determined experimental endpoint, aligning with the principles of the 3Rs (Replacement, Reduction, and Refinement) for improving animal welfare. In some cases, this limit exceeded the last day of measurement, and the mice were immediately euthanized. A sample size of n ≥ 4 per group was used throughout to achieve a statistical significance of P < 0.05. Mice were randomized into treatment groups. Blinding was performed. All tumors were included for analysis. Differences in survival were determined using the Kaplan–Meier method, and the P-value was calculated by the log-rank (Mantel-Cox) test.

### Reagents

Recombinant mouse IFNγ (PMC4031) was purchased from Gibco, mouse TNF-alpha Recombinant Protein (RMTNFAI) was purchased from Thermo Fisher Scientific, etinostat (MS-275) (S1053-SEL-10mg) was purchased from Stratech Scientific. All compounds were dissolved in DMSO (0.1% final concentration). All reagents were used at the indicated concentrations.

### Cell proliferation

Cells were fixed and stained with 0.5 %w/v crystal violet (Sigma) in 4 %w/v paraformaldehyde (PFA) in phosphate-buffered saline (PBS) for at least 30 minutes. Fixed cells were solubilized in 2 %w/v sodium dodecyl sulphate (SDS) in PBS, and absorbance was measured at 595 nm using a Biotek Synergy™ H1 Hybrid Multi-Mode Reader.

### Gene silencing

For siRNA-mediated silencing of Hdac2, cells were seeded in 6-well plates (1 × 10^5^ cells/well), 12-well plates (3.6 x 10^4 cells/well), and 24-well plates (2 x 10^4 cells/well), and incubated overnight. The next day, cells were transfected with siRNAs using DharmaFECT 1 Transfection Reagent (Horizon Discovery) according to the manufacturer’s guidelines. After 24 – 60 hours of incubation with the transfection mixture, the cell culture medium was replaced, and cells were incubated for 1–3 days at 37 °C.

### RNA sequencing

For M4-B2905 cells treated with siHdac2, HDACi, or TNFα, total RNA was isolated from triplicate samples using the RNAeasy mini kit. RNA integrity was assessed on an Agilent 2200 TapeStation (Agilent Technologies). RNA samples (∼1 μg) were submitted for RNA sequencing (100 nt paired-end reads, <30 million reads per sample) using an Illumina HiSeq4000. Three samples per condition were sequenced. Sequence data was collected using the Hiseq Software Suite (version 3.4.0). Read quality was assessed using FastQC. Raw reads were trimmed using trimmomatic (version 0.36.6; sliding window trimming with 4 bases averaging and average quality minimum set to 20). Trimmed reads were aligned to the reference genomes hg38_analysisSet (human) or mm10 (mouse) using HISAT2 (version 2.1.0; default parameters). Aligned reads were counted against GENCODE release 25 (human) or GENCODE release M14 (mouse) using htseq-count (version 0.9.1).

### Western blot

Total proteins were extracted using SDS lysis buffer (4 %w/v SDS; 20 %v/v glycerol; 0.004 %w/v bromophenol blue; 0.125 M Tris-Cl, pH 6.8; 10 %v/v 2-mercaptoethanol) and sonication (50 kHz for 30 s; VibraCell X130PB, Sonics Materials) at 4 °C and subsequently denatured at 95 °C for 5 min. Proteins were separated on RunBlue 4-12 %w/v bis-tris polyacrylamide gels (Expedeon) and then transferred onto iBlot PVDF membranes (ThermoFisher) using the Wet/Tank Blotting Systems (Bio-Rad). Membranes were probed overnight at 4 °C in blocking solution containing the primary antibody. Primary antibodies used in this work were HDAC2 (3F3) (Cell Signaling Technology, 5113S), GAPDH (Proteintech, 60004-1-Ig), and Cleaved Caspase-8 (Asp387) (Cell Signaling Technology, 9429S). This was followed by incubation with the appropriate secondary antibody for 1.5 h at room temperature. Signals were developed using the Clarity Max Western ECL blotting substrate (Bio-Rad) and acquired on a Gel Doc XR+ Gel Documentation System (Bio-Rad). Images were analyzed using Image Lab™ Software (version 3.0.1.).

### Statistical analysis

Statistical analysis calculations were carried out using Microsoft Excel and GraphPad Prism software 10.1.2. For each of the experiments, the statistical experiment was performed separately. P-values < 0.05 were considered statistically significant. All data were expressed in the form of mean ± SEM error unless otherwise specified.

### Multiplex staining of tumor sections

Tumor sections were stained with Picrosirius Red and Fast green to obtain the highest quantitative results, then imaged using a slide scanner, both in brightfield and polarized light to visualize collagen fibres[48]. Tumor sections were stained for a panel of antibodies that includes F4/80 (Cell Signaling 24358S) and CD3 (Cell Signaling 47865S) using Cell Signaling SignalStar Multiplex IHC in Leica BOND according to manufacturer instructions. Briefly, following dewaxing, melanin bleaching (H_2_O_2_ 4% 60°C, 30 min) and antigen retrieval (EDTA), antibodies were added to the slide. Antibodies were conjugated to their unique fluorescent complementary oligos, amplified, and imaged at the bioimaging facility using an Olympus VS200 slide scanner using the 20x objective. Alternatively, slides were immunostained with Sox10 (Abcam ab310390 AF594), Rab38 (Proteintech 12234-1-AP, secondary Invitrogen A-21245 AF647), and DAPI using Leica BOND, then scanned at the appropriate fluorescence channels as above.

### Capsule analysis using Picrosirius stain under polarised light

Polarized light images were analyzed using QuPath scripts[49]. The input region was first down-sampled to 4 μm/px for less computationally expensive resolution. The workflow then branches into two parallel paths. Script 1 – Collagen Segmentation. Detects collagen/tumor regions via color-based masking, followed by blurring, dilation, blob detection, and union of geometries to form annotated regions. Script 2 – Capsule Segmentation. Identifies capsule interiors by masking dark regions while including user-defined ignore* areas, then applies blurring, erosion, and blob detection. The capsule void is rendered as a filled red region, and its surrounding ring is computed through dilation and drawn in blue. Script 3 – Full Tumor. The geometric union of S1 and S2 is dilated and refined through a final blob detection to define the final annotated region.

### CD3+ cell quantification in capsule regions

As SignalStar staining for CD3+ cells (see above) was done on adjacent sections to the picrosirius stain, Qupath annotations (Capsule, Capsule Edge, Whole tumor) produced using capsule analysis scripts were manually copied, rotated, and scaled from the picrosirius section to fit the SignalStar tumor section. CD3+ cells were quantified in each capsule and its matching edge, the proportion of CD3+ cells out of the entire cell population of each region was extracted, then the fold change of each capsule edge to its capsule was calculated, per tumor, per treatment, and knockdown. A Wilcoxon test was used to determine whether the fold change difference per group was different from the null hypothesis (FC=1) (using R).

### Processing of embryonic Dct+ neural crest data

Reads were aligned to a custom CellRanger (v6.1.0) reference generated using the mm10 genome assembly with the sequence of the eGFP construct added as a separate chromosome. Cells with more than 15% of reads aligning to mitochondrial genes were discarded. We ran SoupX on each embryonic sample to remove any contamination from ambient RNA in each cell[50]. For each embryonic time-point, read counts were normalized using the NormalizeData function in Seurat[51] (v4.0.1) with default parameters. The top 10000 most variable features were chosen to carry out PCA, and the first 50 PCs were used to cluster cells using the FindNeighbors function (at a resolution of 0.1) and to generate a UMAP embedding of cells. DoubletFinder[52] was used (with parameters pN = 0.25 and the first 50 PCs) to detect putative doublet cells that were then removed from further analyses. To derive regulons, we ran SCENIC[53] with its default parameter settings. We normalized single-cell regulon activity scores as described in the section on gene set activity normalization and retained regulons with a mean normalized activity score of 0.1 for further analyses. To derive cluster-specific regulons, we averaged normalized gene set activity scores across cells in each cluster and z-scored the mean activity scores across clusters. The top ten differentially active regulons (with a mean activity z-score > 1) in each cluster were chosen as cluster-specific regulons. We next filtered our data based on summed Sox10 and Sox9 expression to confirm that cells were genuinely of the NCC lineage. Summed expression values of zero were excluded. Excluded cell clusters included an Immune subpopulation of E15.5 cells that expressed the macrophage marker genes, Cx3cr1, and Mrc1. Two mesenchymal-like clusters, expressing matrix-remodeling genes, were also excluded. To back up this decision, we confirmed that none of the excluded clusters overlapped pre-existing neural crest single cell data developed by Kastriti et al. (Supplementary Fig. 2c) and were not co-enriched with any Sox10+ clusters in the Kastriti et. al data (Supplementary Fig. 2d). Of note, we could detect GFP RNA in all excluded clusters (Supplementary Fig. 2c), yet the lack of neural crest Sox gene expression suggests these cells may have leaky expression of the transgene.

### Demultiplexing embryos in embryonic melanoblast data based on germline variants

To check if each cluster contained cells from multiple embryos, we used cellSNP[54] to detect FVB-specific SNPs amongst expressed reads in each cell based on the FVB GRCm38 germline variant database, dbSNP v142, (https://ftp.ebi.ac.uk/pub/databases/mousegenomes/REL-1505-SNPs_Indels/strain_specific_vcfs). We then ran Vireo[55] with the number of donors set to three for E11.5 and four for E15.5 datasets. Vireo uses the pattern of variants in each cell to infer which embryo was its source and labels cells as “unassigned” if its statistical model cannot resolve the source embryo for a cell. With default settings, Vireo was able to assign 49.3% of E11.5 cells and 24.7% of E15.5 cells to a single embryo.

### Integration and co-clustering analysis of Dct+ neural crest and published Sox10+ embryonic neural crest

The SMART-seq2-derived read count matrix of Sox10+ neural crest data from mouse embryos[15] , along with associated cell annotations, was downloaded from the GEO database (GSE201257). Read counts were normalized with Seurat (v4.3.0) using log-normalization in the NormalizeData function. Batch correction and integration with our embryonic data were performed with the IntegrateData function in Seurat based on the top 5000 most variable genes and 50 PCs. This procedure was performed separately with Mes.1 and Mes.3 cells from E15.5 being either included or excluded. The integrated Dct+ and Sox10+ neural crest data were clustered at a resolution of 0.8 using the FindClusters function in Seurat based on the expression of 5000 genes selected by Seurat post-integration. We used Fisher’s exact test to check if each post-integration cluster was enriched for an annotated cell type in the Dct+ or Sox10+ neural crest data. We then assembled a co-enrichment network by drawing an edge between a Dct+ and Sox10+ annotated cell type if they were both enriched in at least one post-integration cluster with an adjusted p-value < 0.1.

### Derivation of developmental gene modules

We merged cells across E11.5 and E15.5 time-points and inferred differentially expressed genes for each cluster using the FindAllMarkers function in Seurat, which returned genes with an average log2-fold change of at least 0.25. The MAST test was used to estimate p-values with the cell cycle score of each cell being considered as a covariate. Genes with an adjusted p-value less than 0.1 were retained. Amongst the up-regulated genes in each cluster C, we construct a gene set G_C_ by discarding all cell cycle genes along with any genes whose expression has a Spearman correlation coefficient of at least 0.6 with cell cycle genes. We then performed a lineage-based correlation filtering to retain co-expressed genes within each cluster C as follows. We grouped clusters into four lineages: SCP – SCP.1, SCP.2, SCP.Early, Mes – Mes-1, Mes-2, Mes-3, Mes-Early, Mel – Mel, Mel-Early, Neural – NPC2, Sensory (immune cluster is not considered for further analysis). We do not group NPC1 and Notochord DGMs into a single lineage owing to their mapping to multiple developmental lineages in Sox10+ neural crest data. For a given cluster C belonging to a lineage L, we compute the co-expression score (Spearman correlation) between every gene pair in G_C_ across cells in C and repeat this process for every other cluster C’. If a gene pair is correlated across clusters in L as well as in other lineages, then that gene pair is discarded. Amongst genes in the retained gene pairs, we retain at most 200 genes (based on their average log2-fold change) as the DGM of that cluster.

### Computing gene set activity scores in bulk and single-cell RNA-seq data

Single-cell RNA-seq data: We used AUCell[53] to compute raw gene set activity scores for a given collection of gene sets (DGMs or otherwise). AUCell computes a gene set activity score based purely on the gene expression of that cell and does not depend on the gene expression profile of other cells. AUCell also computes multiple enrichment thresholds at which a gene set can be considered active. We use the Global_k1 threshold and label cells whose AUCell score for a given gene set is above this threshold to be enriched for that gene set.

Mapping hybrid states in single-cell RNA-seq data: We score the activity of DGMs in all cells in a single-cell cohort (across sample donors) using AUCell. A DGM is enriched in a cell if its AUCell activity score exceeds its Global_k1 threshold. We consider a cell to belong to a given lineage L if the cell is enriched for the activity of at least one DGM representing L. If a cell is enriched in DGMs of more than one lineage, we consider that cell to be in a hybrid state.

Bulk RNA-seq data: To score gene set activity in bulk RNA-seq data for a given cohort, count data is transformed into a log-TPM scale. The mean log-TPM score of all foreground gene sets is first computed. Each gene in the collection is assigned to one of 20 bins based on its mean expression. A set of 100 control gene sets was constructed, consisting of genes from the union of all gene sets as described above. The mean expression of each control gene set is computed, and the 95^th^ percentile is subtracted from the activity of the foreground gene set to obtain the normalized activity score.

### TCGA analysis

Log-normalized FPKM values for melanoma tumor RNA-seq samples (TCGA code SKCM) were obtained from the Toil recompute project on the UCSC Xena platform. Tumor purity scores were obtained from the TCGA PanCan Atlas Supplemental Data collection. TIDE scores[56] for each sample were computed from the TIDE website (http://tide.dfci.harvard.edu/). DGM activities were evaluated across the TCGA SKCM cohort as described above. TIDE estimated whether each tumor sample contained TILs, based on which we grouped samples into TIL+ and TIL-groups. To determine if a DGM is differentially active in TIL+ samples compared to TIL-samples, we used the glm function in R to fit a linear model of DGM activity versus TIL status and the TIDE-computed CAF (cancer-associated fibroblast) score of each sample.

### Immune checkpoint blockade analysis

Re-processed read counts of Van Allen et al.[57], Liu et al.[58], Gide et al.[59], and Riaz et al.[60] were obtained from reprocessed immunotherapy cohorts. Read counts for Van Allen et al. were obtained from the source publication[57]. Annotated single-cell RNA-seq data of malignant cells (Pozniak et al.) were provided by the authors[29]. In all analyses, only RNA-seq samples of pre-treatment biopsies of patients receiving either anti-PD1 or anti-CTLA4 were analyzed. A Cox mixed-effects regression was carried out to determine the hazard ratios of DGM activity on patient survival, with the patient cohort label of each sample used as a mixed effect to account for cohort-level differences in survival.

### Analysis of Vallius et al. microregion data from early melanoma lesions

DGM activities were computed in RNA-seq from microregions as described above. The R lme4 function from the lmerTest package was used to compute t-values associated with DGM activity differences in P vs N, MIS vs P, RGP vs MIS, and VGP vs RGP contrasts (with patient ID as a random effect to account for multiple microregions coming from each patient). In the analysis of DGM activity changes due to TILs, the lme4 function was used to estimate the significance of t-values of the brisk TIL vs no-immune and non-brisk TIL vs no-immune contrasts (with patient ID set as a random effect).

### Estimating the background distribution of hybrid states in M4-BRN2 cells

We used Gaussian copula sampling to simulate DGM activities for a given scRNA-seq dataset. Gaussian copula sampling involves first drawing samples from a multivariate normal distribution which requires two sets of inputs – the mean of each random variable, along with the covariance matrix that captures correlations between all pairs of random variables. The background covariance matrix we used was obtained by evaluating DGM activities in our iDct-GFP+ neural crest dataset and computing covariances between all pairs of DGM activities. This matrix represents the expected correlation between any pair of DGM activities when DGMs are evaluated in any scRNA-seq dataset and represents lineage relationships between Mel, Mes, Neural, and SCP lineages in the embryo.

After computing this background matrix, we then used AUCell to compute DGM activities in M4-BRN2 cells and assign each cell to a lineage based on whether a DGM activity is higher than its respective threshold T. We then draw a sample matrix *X*_*N*×*d*_ consisting of *N d*-dimensional vectors (where *N* is the number of M4-BRN2 cells and *d* is the number of DGMs) of DGM activities from a multivariate Gaussian distribution whose means are the average activity of each DGM amongst M4 cells and whose covariance matrix is the one computed above from iDct-GFP neural crest data. Let *G*_1_, *G*_2_, … , *G_d_* represent the marginal Gaussian cumulative distribution function of each DGM’s sampled activities and *F*_1_, *F*_2_, … , *F_d_* represent the empirical cumulative distribution function of each DGM’s observed activities in M4-BRN2 cells. We then transform each entry *X_ij_* into *Y_ij_* = *F*^−1^ (*G_j_* (*X_ij_*)) where *G_J_*(*X_ij_*) follows a *U*[0,1] distribution and *F*^−1^ (*G_j_* (*X_ij_*)) follows the distribution specified by *F_j_*. We then determine if *Y_ij_* > *T_J_*, where *T_J_* is the enrichment threshold for DGM j computed amongst M4-BRN2 cells, to determine if the j’th DGM is active in cell i and assign the respective lineage(s) to the cell. The fraction of cells in each hybrid state is then stored.

The above sampling procedure is repeated a hundred times to build a distribution of the expected fraction of cells in each hybrid state. A two-sided p-value is computed for the observed fraction of cells in each hybrid state after fitting a normal distribution to the background fractions simulated above.

### Processing of metastatic single-cell RNA-seq data

Reads were aligned to a custom CellRanger (v6.1.0) reference generated using the hg38 human genome assembly with the sequence of the eGFP construct added as a separate chromosome. Cells with more than 15% of reads aligning to mitochondrial genes were discarded. For each metastatic time-point, read counts were normalized using the NormalizeData function in Seurat[51] (v4.0.1) with default parameters. DoubletFinder[51] was used (with parameters pN = 0.25 and the first 30 PCs) to detect putative doublet cells that were then removed from further analyses. The SCTransform method[61] was used to perform integration and batch correction (based on the first 20 PCs with k.score = 20 and k.weight = 20). Clustering was performed on the integrated data with a resolution of 0.2 and based on the first 30 PCs.

We performed two steps of computation to separate malignant from non-malignant cells. First, we ran inferCNV (inferCNV of the Trinity CTAT Project) to infer the number of gene-level deletions and amplifications of cells in each cluster using droplet scRNA-seq data from the Tabula Muris project[62] as a reference background dataset. We then scored each cluster using AUCell with signature genes of every cell type from the healthy lung background data. We counted the number of gene-level CNVs (amplifications and deletions) based on whether the CNV score assigned to the gene was above the 97.5^th^ percentile (amplification) or below the 2.5^th^ percentile (deletion) of CNV scores amongst background cells. Clusters that had a high CNV burden and did not score highly on markers of healthy lung cell types were declared as malignant cells.

Out of 9 cell clusters, one malignant cluster was excluded for being enriched in apoptotic signatures. Another cluster expressed macrophage marker genes and had an intermediate CNV score, potentially representing macrophages that have engulfed a malignant cell. One intriguing cluster expressed endothelial marker genes and had an intermediate CNV score. This cluster potentially contains both metastatic and normal cells with an endothelial phenotype or may represent metastatic cells that have transdifferentiated to an endothelial subtype, which has been noted in another melanoma metastasis study[63]. While this is a notable feature of this model, it did not appear to be linked to the co-option of melanocytic lineage biology and was not pursued further.

### SCENIC analysis to determine upstream regulators of melanoma NS-Hybrid State

SCENIC was run with default options on all cells in our embryonic Dct+ NCC dataset to derive regulons for each transcription factor. The target genes of each inferred regulon were then scored across malignant cells in both the M4-BRN2 mouse metastatic single-cell RNA-seq dataset and the Jerby-Arnon et al. patient cohort using AUCell. Within the M4-BRN2 dataset, we fit a linear mixed model of each cell’s regulon AUCell score versus its hybrid state label (NS-Hybrid, Other-Hybrid, or Singleton) and the logarithm of total reads per cell as fixed effects and the time-point from which the cell was sampled as a random effect. The same fixed effects were used to regress each regulon’s AUCell score in the Jerby-Arnon et al. cohort, but the patient ID was used as a random effect. The t-value associated with the contrast NS-Hybrid minus Singleton was stored in each regression, where the TFs associated with regulons amongst the ten highest t-values were considered for experimental validation.

### Analysis of bulk RNA-seq from siHdac2 and HDACi treatments on M4 and YUMM2.1 cell lines

Reads were aligned to the mm10 genome using the salmon pseudoaligner and the salmon index constructed using Gencode vM9 transcript annotation. DGM activities were computed based on log-TPM normalized expression values of each gene. DESeq2 was used to estimate log-fold changes of expression between siHdac2 and siControl conditions. FGSEA was used to estimate enrichment scores of mouse orthologs of gene sets of the Hallmark pathways and Reactome extracellular matrix pathways sourced from MsigDB.

## Acknowledgements

This research was supported [in part] by the Intramural Research Program of the National Institutes of Health (NIH). The contributions of the NIH author(s) were made as part of their official duties as NIH federal employees, are in compliance with agency policy requirements, and are considered Works of the United States Government. However, the findings and conclusions presented in this paper are those of the author(s) and do not necessarily reflect the views of the NIH or the U.S. Department of Health and Human Services. This work was supported by U.S. National Cancer Institute grants (ZIABC012176-01) to Sridhar Hannenhalli and (ZIABC008756-36) to Glenn Merlino.

Components of this study were funded by the British Skin Foundation Young Investigator Award, 023/YI/22. This work was supported by Cancer Research UK [RCCCDF-Nov24/100004], The Wellcome Trust [204796/Z/16/Z], The University of Manchester FBMH Dean’s Prize award, and NCI Director’s Innovation award, all awarded to KLM. Components of this study were funded by grant 821034 from the Melanoma Research Alliance (MRA) and grant 22-0091 from Worldwide Cancer Research (WCR) awarded to AH. TV is supported by a Research Scholar Grant, PF-24-1316850-01-CD, from the American Cancer Society. Melanoma research in the Sorger lab (TV and YS) is supported by the Harvard Ludwig Center and the Mark Foundation for Cancer Research. Cell sorting and flow cytometry analyses were performed at the NIH, NCI LGI Flow Cytometry Core, supported by funds from the Center for Cancer Research (CCR). We acknowledge support from the CCR Single Cell Analysis Facility for single cell RNA-Seq captures, library preparation, sequencing, and primary data processing was funded by FNLCR Contract 75N91019D00024. This work utilized the computational resources of the NIH HPC Biowulf cluster (http://hpc.nih.gov). We thank Histoserv, Inc. (Germantown, USA) for expert advice and histology and immunolabeling of samples (particularly Pragun Vohra). We thank the Slide scanning service, NCI (in particular Jennifer Dwyer), Biological Services Facility (UoM), Sequencing core (UoM), and Bioimaging (particularly Peter March and James Bagnall). For useful conversations, we thank: Marcus Preedy, Mike White, Kieran Sefton, Martin Lowe, Joan Chang, Andrew MacDonald, Michael Smith, William J Pavan, and Richard White. We thank Peter Sorger for helpful comments. We are grateful to Kun Wang for reprocessing RNA-seq data from ICB cohorts and to Maxwell Lee and Howard Yang (NIH) for early data processing. We thank Rachel Belote for annotated single-cell RNA-seq data of human melanocytes and Joanna Pozniak for annotated single-cell RNA-seq data of human melanoma patient tumors. For technical help assisting tissue harvesting and tumor growth studies, we thank Wei Zhang, Christopher Jones, Jiaqing Lang, and Sorayut Chattrakarn (UoM). We used OpenAI’s ChatGPT (accessed May 2025) to assist with summarizing results and editing manuscript text.

## Author Contributions

KLM, VG, GM, SH, CPD, and AH conceptualized the study. KLM, VG, CWW, RL, LO, EPG, EW, and MH developed methodology.; VG, LO designed software. KLM, VG, RL, and CWW performed validation experiments. VG, KLM, CWW, RL, LO, FL, YS, and RMS performed formal analyses. VG, CWW, RL, KLM, YJ, EW, AS, MAC, SC, JE, and CS conducted research investigation. RPG, TV, YS provided novel resources, VG, RL, and LO performed data curation. KLM, VG, AH wrote the original draft, KLM, VG, AH, CPD, TV, YS, CWW, RL, SH, GM reviewed and edited the manuscript. VG, KLM, RL, CWW, LO, FL, RMS prepared figures for published work (visualization). KLM, GM, SH, AH, RL, VG undertook supervision roles. KLM, VG managed and coordinated project administration. KLM, GM, SH, and AH provided funding acquisition.

## Data Availability

Unpublished data and code will be made available upon request. Once published, all data and code will be made freely available on an online repository.

