## Supplementary Figures and Table for "Transitory Schwann Cell Precursor and hybrid states underpin melanoma therapy resistance and metastasis"

### Supplementary Figure 1

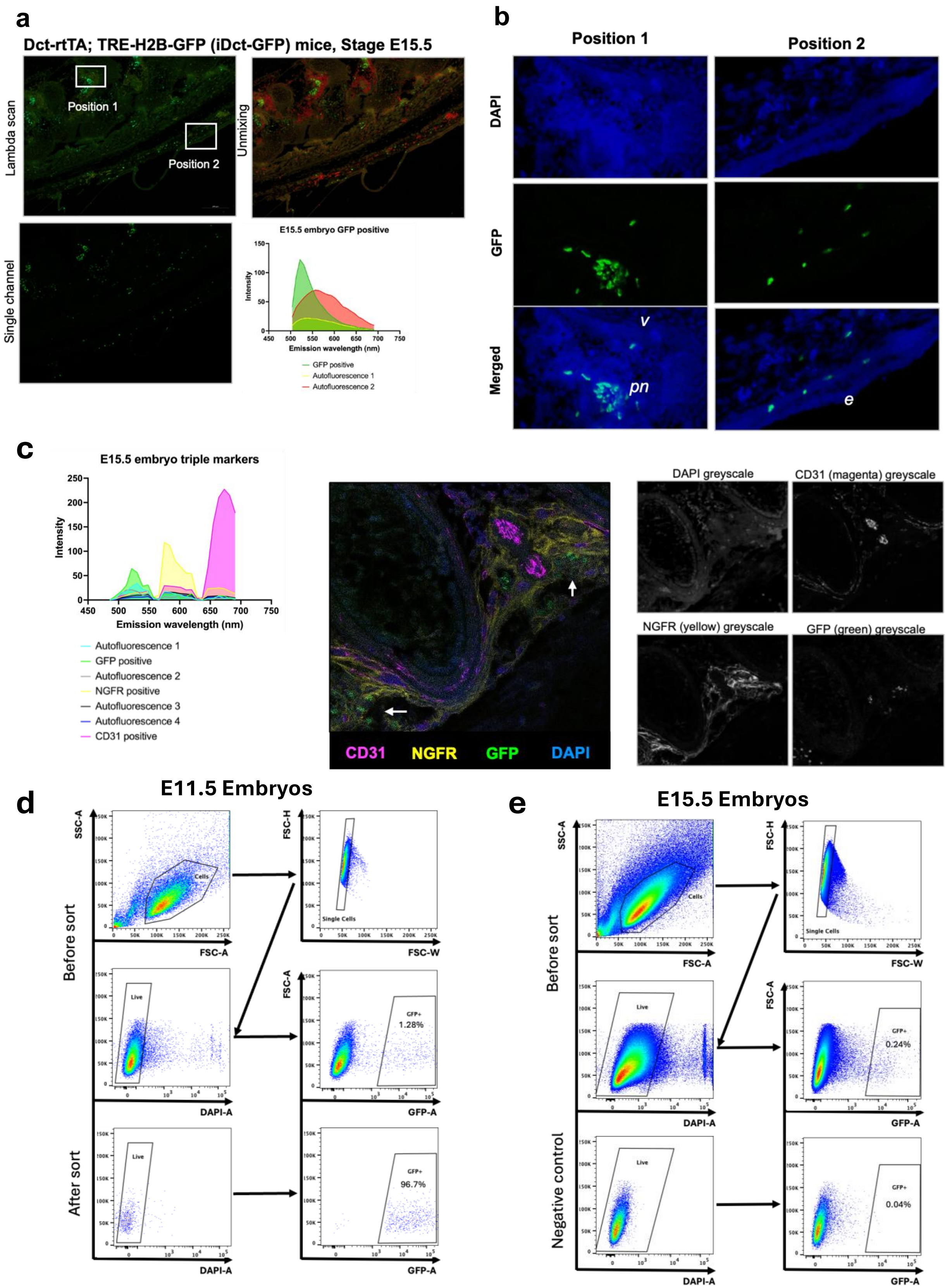

**Supplementary Figure 1. Visualization and capture of GFP+ NCCs from the iDct-GFP transgenic mouse.** (**a**-**b**) Anti-GFP immunofluorescence of paraffin-embedded Embryonic day (E) 15.5 embryo samples. Sagittal sections of double allele Dct-rtTA: TRE-H2B-GFP (iDct-GFP) mice. The dorsal (back) region of the embryo's trunk, including skin (epidermis, e) and intercostal bundles of GFP+ cells, likely surrounding nerves (peripheral nerves, pn). The more ventral region of the image indicated, v. Lambda scans (a) using confocal microscopy and spectral unmixing allow accurate identification of GFP-specific emission spectra. (**c**) Immunofluorescence of E15.5 sagittal sections with results from Lambda scan and spectral unmixing. Left: Intensity plot of specific emission spectra. Middle, Right: Anti-CD31 (blood vessels) shown in magenta, anti-NGFR (Neural Growth Factor Receptor) shown in yellow, anti-GFP (GFP+ cells) shown in green, and DAPI in blue. (**d**-**e**) Fluorescence-Activated Cell Sorting gating strategy to capture GFP+ NCCs in iDct-GFP (Dct-rtTA: TRE-H2B-GFP) mice compared to TRE-H2B-GFP one allele control mice. Pregnant mice were fed doxycycline chow from conception. DAPI was used to distinguish live cells from dead cells.

### Supplementary Table 1

| Sample Name | Estimated Number of Cells | Mean Reads per Cell | Median Genes per Cell | Sequencing Saturation | Total Genes Detected | Median UMI Counts per Cell |
|---|---|---|---|---|---|---|
| E15.5 | 4,230 | 76,980 | 3,702 | 65.80% | 21,091 | 12,487 |
| E11.5 | 5,262 | 71,485 | 4,138 | 50.80% | 20,679 | 18,319 |

### Supplementary Figure 2

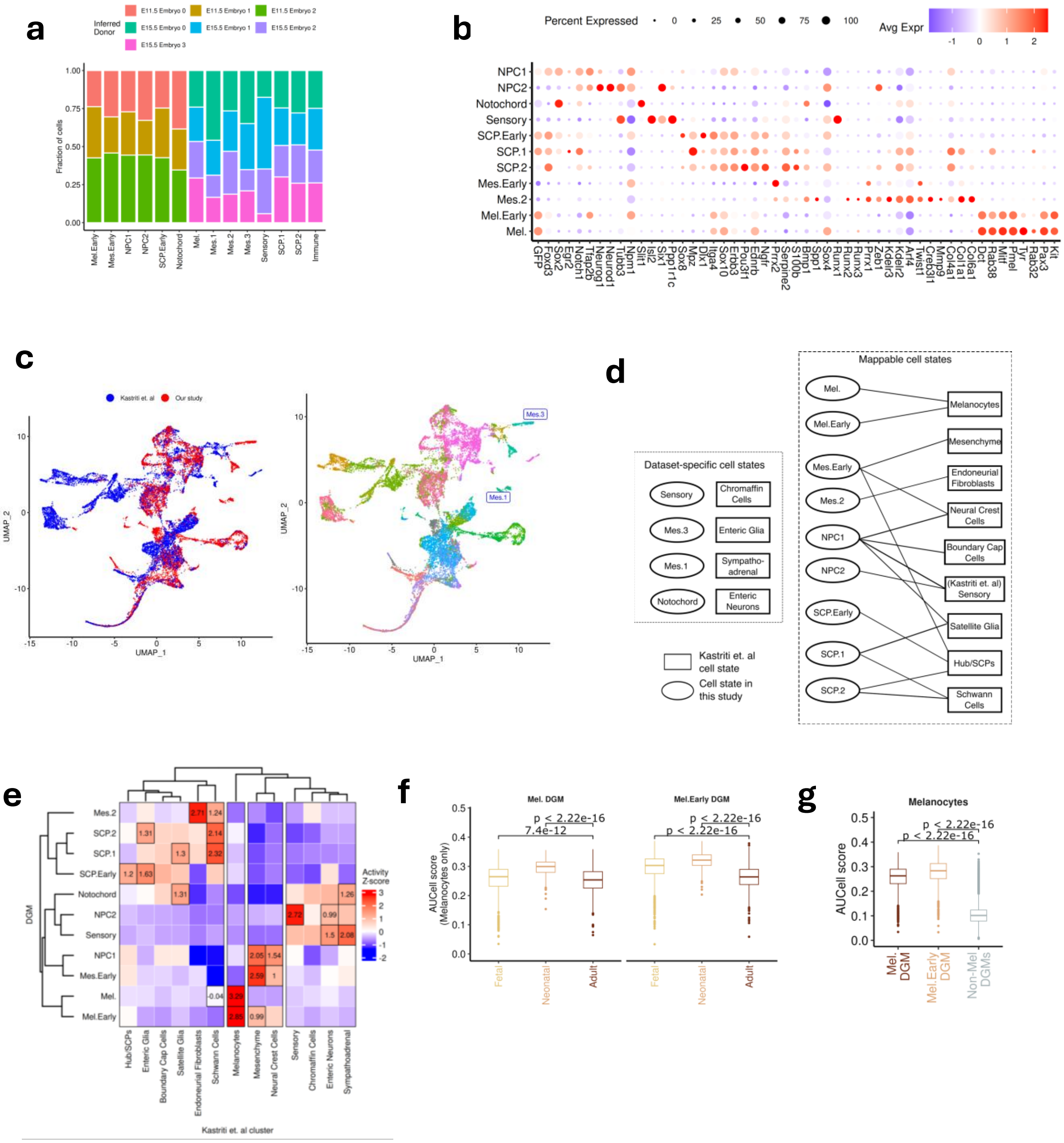

**Supplementary Figure 2. scRNA-seq of GFP+ NCCs from iDct-GFP transgenic mouse. (a)** Contribution of different embryos to cells from each cluster in Dct+ neural crest single-cell RNA-seq data. **(b)** Dot plot visualization of the relative expression of selected neural crest markers in our scRNA-seq data. **(c)** Co-embedding of scRNA-seq of GFP+ NCCs from our study with Sox10+ NCCs from Kastriti et al. Mes.1 and Mes.3 clusters from our study are highlighted. **(d)** Co-clustering network of GFP+ NCC cell states (circles) from our study and Sox10+ NCC cell states (rectangles) from Kastriti et al. Edges indicate clusters from both datasets that significantly co-cluster with each other (adjusted $p < 0.05$, permutation test). (**e**) Hierarchically clustered heatmap of DGM scores across NCCs in Kastriti et al. (**f-g**). Box and whisker plots of DGM scores across melanocytes in Belote et al. Kit+/CD34- scRNAseq data, p-values indicated, two-sided Wilcoxon test. The line represents the median; the box represents the interquartile range**.**

### Supplementary Figure 3

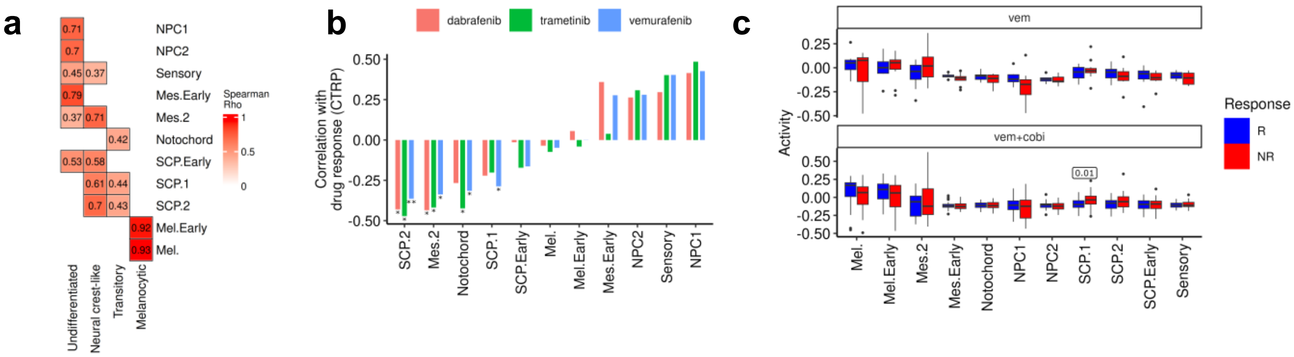

**Supplementary Figure 3. Correlations between DGM expression and clinical phenotypes.** (**a**) Spearman correlation coefficient between DGM scores and Tsoi et al. signature scores evaluated in bulk RNA-seq of Tsoi et al. cell line panel. Only statistically significant positive correlations (adjusted $p < 0.05$, permutation test) are shown. (**b**) Correlation between DGM activity in CCLE cell lines and area above drug response curve to dabrafenib, trametinib, and vemurafenib treatment of CCLE cell lines in the CTRP database. Negative correlation implies that high DGM activity is associated with resistance. ** $p < 0.01$, * $p < 0.05$; two-sided bootstrap test (**c**) DGM scores in bulk RNA-seq of pre-treatment tumor biopsies of responder and non-responder patients receiving BRAFi (vemurafenib) and BRAFi+MEKi (vemurafenib + cobimetinib) in Yan et al. cohort. ** $p < 0.01$, * $p < 0.05$; two-sided Wilcoxon test.

### Supplementary Figure 4

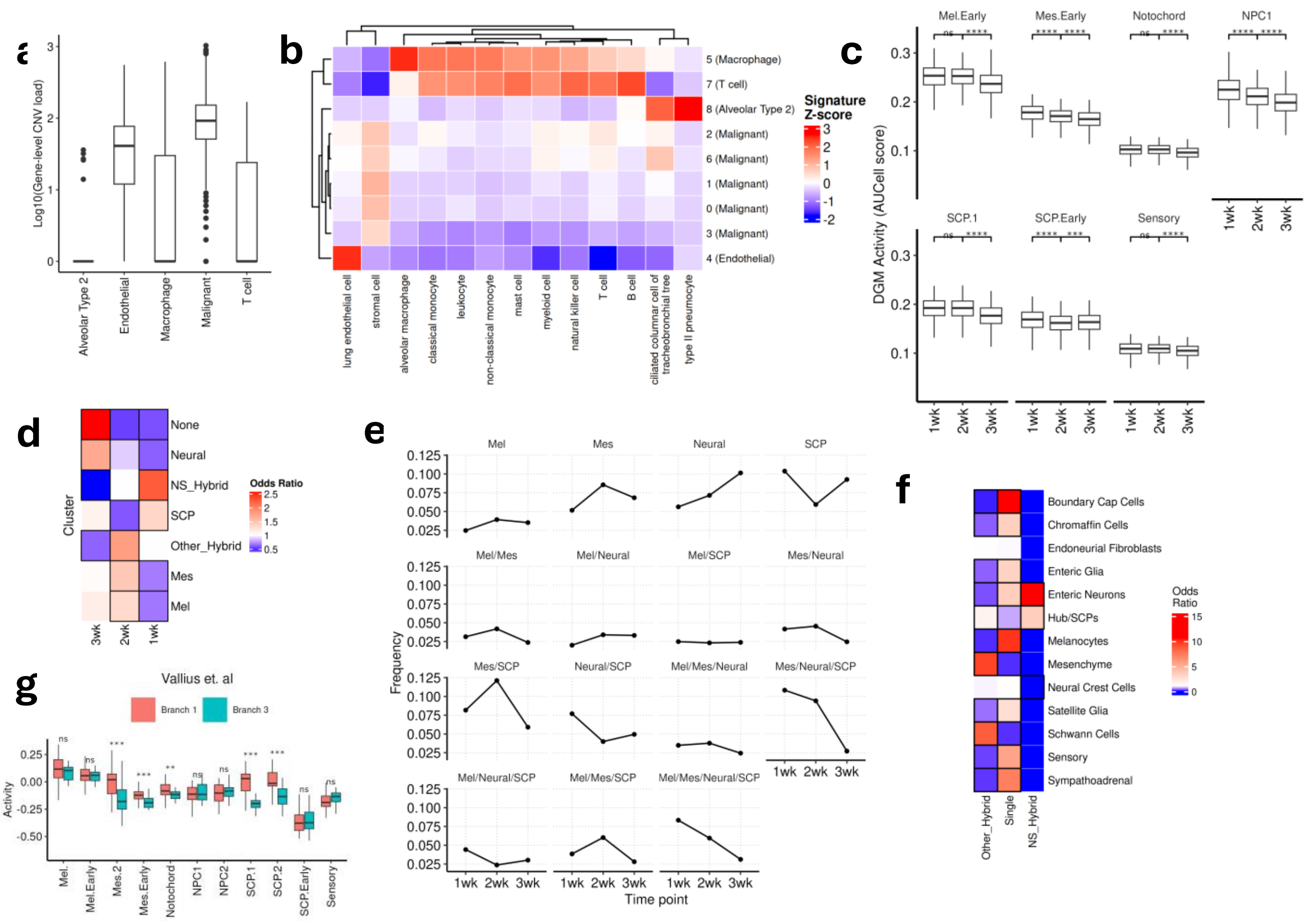

**Supplementary Figure 4. DGM activities in M4-BRN2 single-cell RNA-seq data.** (**a**) Number of gene-level copy number amplifications and deletions in each cell inferred by inferCNV across malignant and non-malignant cells. (**b**) Z-score of AUCell scores of gene sets derived from normal lung populations in lung scRNA-seq data from the Tabula Muris project (columns) across different clusters in M4-BRN2 scRNA-seq data. (**c**) AUCell scores of DGMs evaluated in malignant cells sampled across time-points. **** $p < 10^{-4}$; two-sided Wilcoxon test (**d**) Enrichment of specific hybrid and singleton states across 1-, 2-, and 3-week post-colonization. Squares with black borders indicate significant enrichments (adjusted $p < 0.1$, Fisher's exact test). (**e**) Proportions of malignant cells in singleton and hybrid states across time-points. State labels are at the top of each panel. (**f**) Enrichment of NS-Hybrid, Other-Hybrid, and singleton states across Sox10+ scRNA-seq clusters in Kastriti et al., 2022. Black squares indicate significant enrichments (adjusted $p < 0.1$, Fisher's exact test). (**g**) DGM activity in microregions of early melanoma samples profiled by Vallius et al. and classified as either Branch 1 or Branch 3. **** $p < 10^{-4}$, *** $p < 10^{-3}$, ** $p < 10^{-2}$, * $p < 0.05$; two-sided Wilcoxon test.

### Supplementary Figure 5

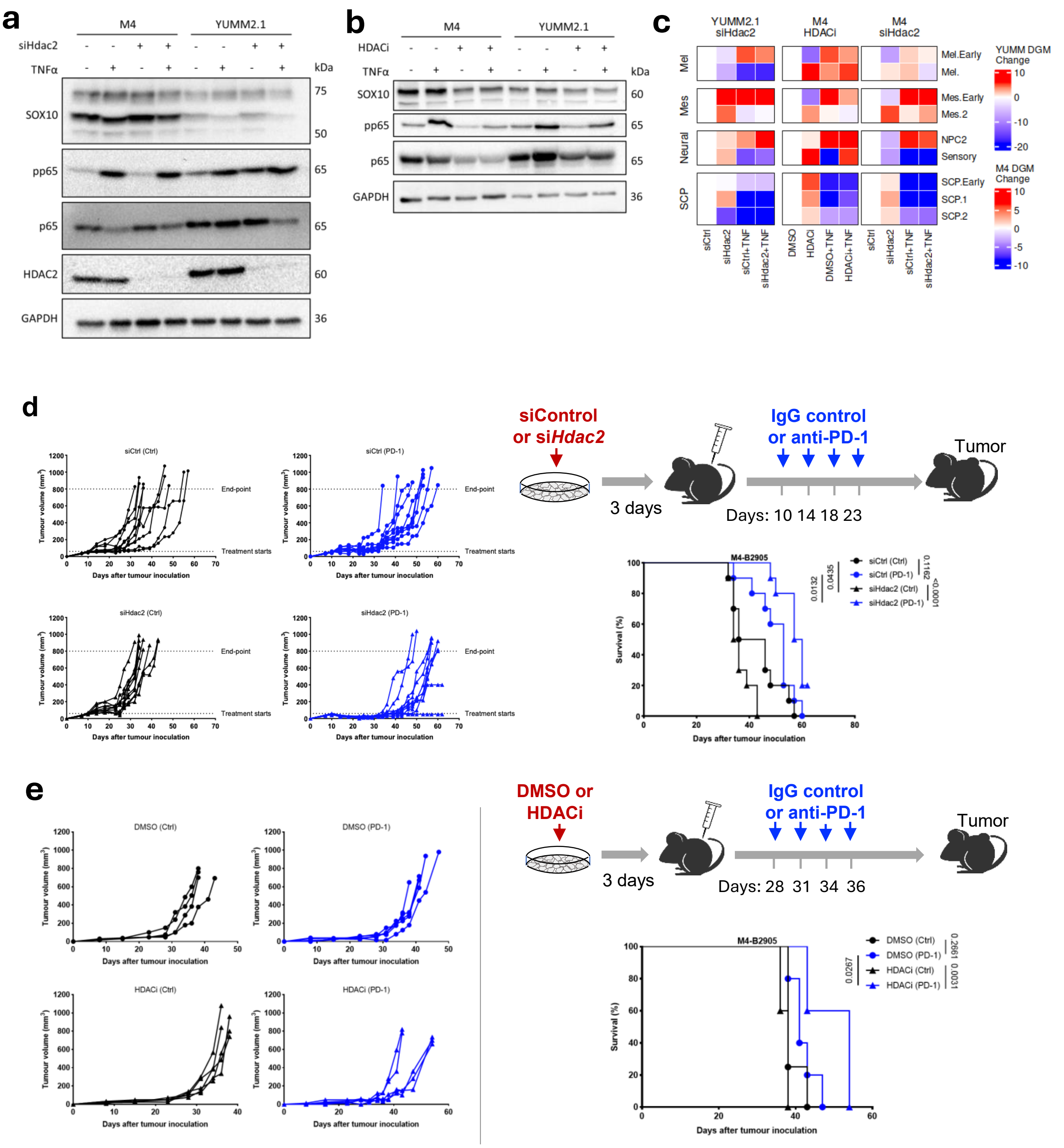

**Supplementary Figure 5. Molecular mechanisms and phenotypes following HDAC loss in melanoma cells.** (**a, b**) Immunoblot staining of cell lysates from B2905 and Yumm2.1 mouse melanoma B2905 cells. (**c**) Relative DGM activity change across Wildtype and *Hdac2* knockdown (siHdac2) (M4) and siHdac2 YUMM2.1 cell lines treated with TNFα or DMSO. DGM activities in DMSO and non-targeting control siRNA are used as baselines for z-score calculation in respective treatment conditions. (**d-e**) Schematic of preclinical experiment design, individual tumor growth charts, and cumulative mouse survival curves for (e) *Hdac2* siRNA knockdown (siHdac2) versus non-targeting control siRNA (siCtrl) or (f) Entinostat (HDACi, HDAC inhibitor) treatment versus DMSO control, treated with IgG control antibody (black) or anti-PD-1 ICI therapy (blue). P-value determined by Kaplan–Meier, log-rank (Mantel-Cox) test.

### Supplementary Figure 6

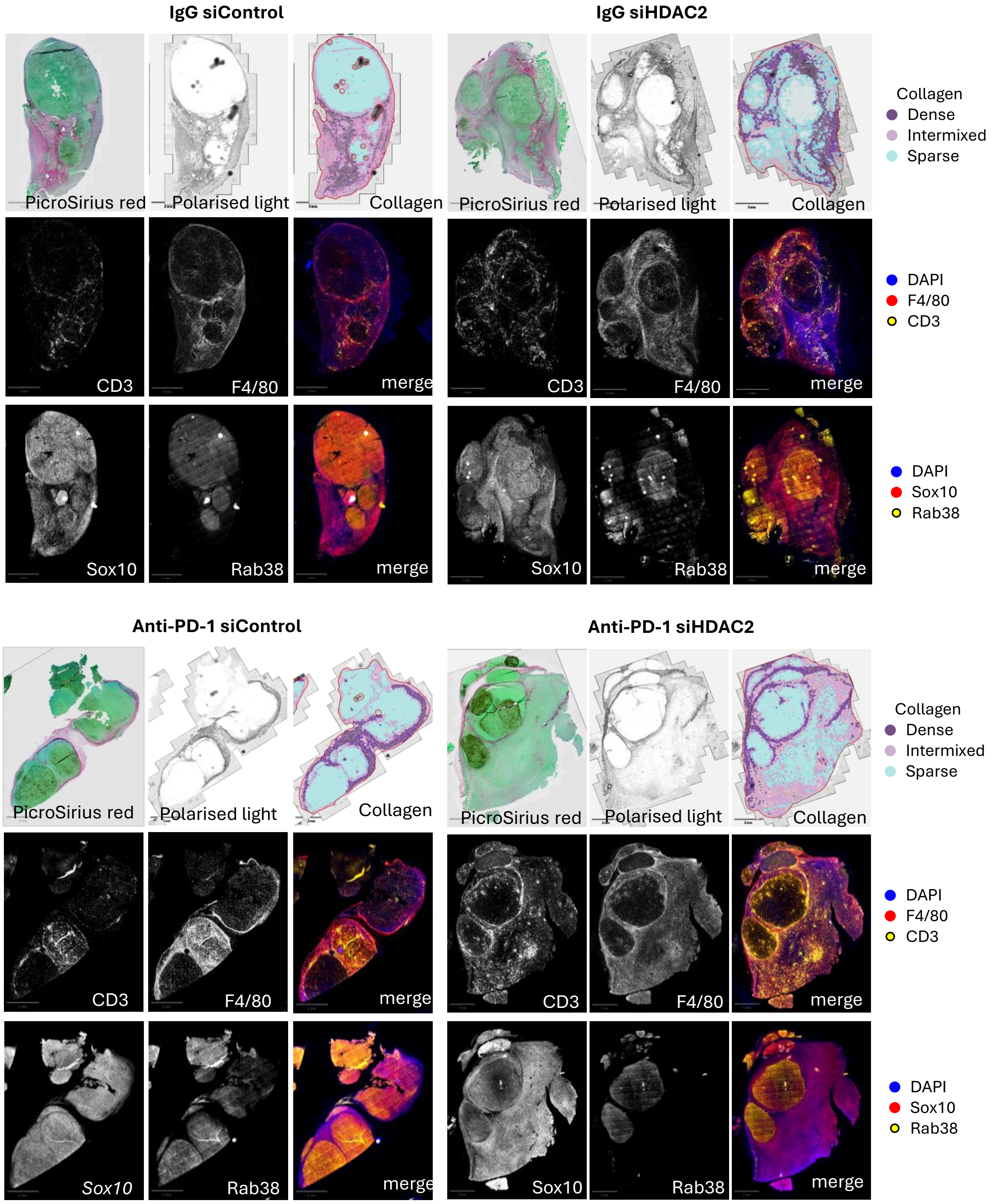

**Supplementary Figure 6. Multiplex imaging of tumors to reveal TME organization.** Multiplex imaging of mouse subcutaneous tumors derived from B2905 cells (Figure 6a-b). Representative example of each treatment modality shown (from n=10 tumors per treatment). Top row (left to right): Brightfield scans of picrosirius red-stained tumor sections, with green counterstain; optical density sum projected image of picrosirius red-stained tumor sections visualized under polarized light; Qupath AI-generated detection of sparse, intermediate, and medium collagen densities based on the optical density sum images. Second row: Immunostaining of tumors with anti-CD3 and anti-F4/80 antibodies. Third row: Immunostaining with anti-RAB38 and anti-SOX10 antibodies.

### Supplementary Figure 7

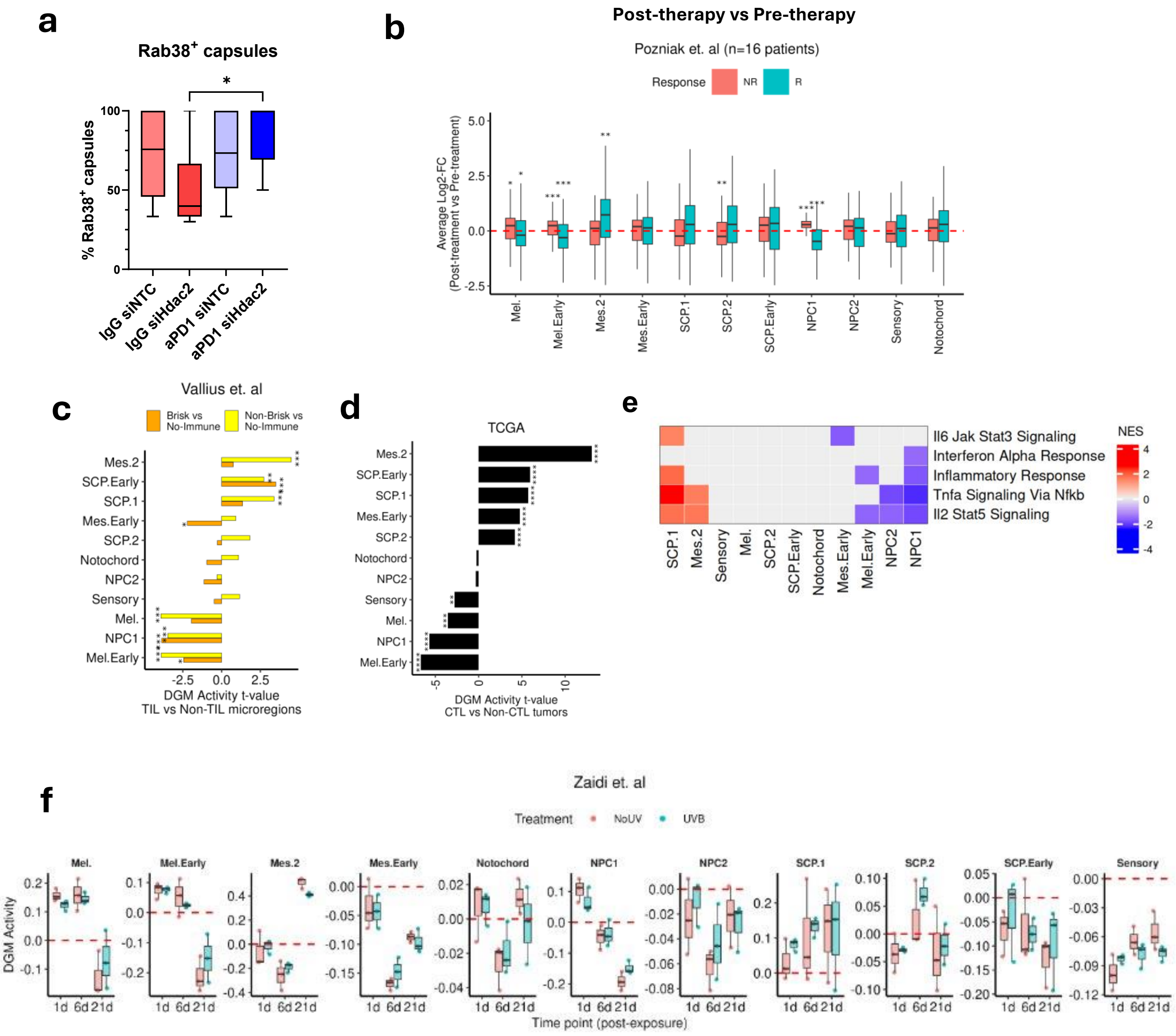

**Supplementary Figure 7. Lineage states and inflammatory changes over time in tumors and normal tissue.** (**a**) Percentage of capsules per tumor with RAB38+ expression. siNTC, Non-targeting control siRNA; si*Hdac2*, *Hdac2*-targeted siRNA; IgG, immunoglobulin control; aPD1, anti-PD-1 treated. Kruskal-Wallis test, Dunn's multiple comparison, * $P<0.05$. (**b**) Average Log2-FC of DGMs in post-therapy vs pre-therapy malignant cells (from Pozniak et al.) in responders and non-responders to anti-PD1 therapy. **** $p < 10^{-4}$, *** $p < 10^{-3}$, ** $p < 10^{-2}$, * $p < 0.05$; two-sided Wilcoxon test (**c**) t-values of DGM activities between microregions in early melanoma samples containing brisk or non-brisk TILs compared to regions without immune infiltration. Patient ID is set as a random effect in the linear model. **** $p < 10^{-4}$, *** $p < 10^{-3}$, ** $p < 10^{-2}$, * $p < 0.05$; t-test. (d) t-values of DGM activities between TCGA samples with and without TILs. **** $p < 10^{-4}$, *** $p < 10^{-3}$, ** $p < 10^{-2}$, * $p < 0.05$; t-test. (**e**) GSEA normalized enrichment scores of selected MsigDB Hallmark pathways in Dct+ neural crest clusters. Squares in gray indicate no significant enrichment for that pathway (adjusted $p > 0.1$, permutation test). (**f**) Activities of DGMs in microarray data (from Zaidi et al., three replicates in each condition) from mouse melanocytes irradiated by UVB or control.
